# The *Mycobacterium tuberculosis* protein *O*-phosphorylation landscape

**DOI:** 10.1101/2022.02.17.480717

**Authors:** Andrew Frando, Vishant Boradia, Marina Gritsenko, Michael-Claude Beltejar, Le Day, David R. Sherman, Shuyi Ma, Jon M. Jacobs, Christoph Grundner

## Abstract

Bacterial phosphosignaling has long been synonymous with the histidine kinases of the two component systems, but many bacteria, including *Mycobacterium tuberculosis* (*Mtb*), also code for Ser/Thr protein kinases (STPKs). STPKs are the main phosphosignaling enzymes in eukaryotes, but the full extent of phosphorylation on protein Ser/Thr and Tyr (*O*-phosphorylation) in bacteria remains unclear. Here, we explored the global signaling capacity of the STPKs in *Mtb*. We generated STPK loss-and gain-of-function strains and measured the resulting *O*-phosphorylation and transcriptional changes. This deep phosphoproteome shows that *O*-phosphorylation in *Mtb* is an underexplored protein modification that affects >70% of the proteome. The substrate-STPK interactions show an extensive interface with the transcriptional machinery, resulting in regulation of gene expression of ∼30% of *Mtb* genes. *Mtb O*-phosphorylation gives rise to an expansive, distributed, and cooperative network of a complexity that has previously only been associated with eukaryotic phosphosignaling networks.

## INTRODUCTION

Bacteria signal through two branches of phosphosignaling with different chemistries of protein modification (1,2). The His kinases of two component systems (TCSs) have long been considered the mainstay of bacterial phosphosignaling, but phosphorylation on protein Ser/Thr and Tyr (*O*-phosphorylation) is emerging as another important bacterial phosphosignaling mechanism catalyzed by the Ser/Thr protein kinases (STPKs). The bacterial STPKs are similar to the primary phosphosignaling enzymes in eukaryotes (3-5) and have been termed “eukaryotic-like”. The pervasive presence of bacterial STPKs throughout the bacterial kingdom (6) as well as in archaea (7) and the presence of archetypical STPK sequences in bacteria, however, all suggest a common origin at the root of the tree of life. *Mycobacterium tuberculosis* (*Mtb*) encodes 11 STPKs, a relatively large number in relation to its genome size, and nearly as many as TCSs (8,9). All but two STPKs are transmembrane receptors that are likely detecting changes in the environment (8). In this way, the *Mtb* STPKs are primed to be a major conduit for turning extracellular signals into bacterial adaptations.

Many studies have defined physiological substrates and functions for several of the *Mtb* STPKs, in particular the essential PknA and PknB, which broadly affect transcription and metabolism (10,11) and have central roles in coordinating replication and cell division (12). These and other studies point to a more dominant role of *O*-phosphorylation in *Mtb* and perhaps other bacteria than is presently recognized. Phosphoproteomic studies provided the first insights into the full extent of *Mtb O*-phosphorylation (13,14). Despite this emerging picture of *Mtb O*-phosphorylation as an important arm of bacterial and especially *Mtb* cell signaling, the full scope of *O*-phosphorylation is not known, and most STPK cellular functions, signaling pathway organization, regulatory mechanisms, and overall signaling network architecture remain to be characterized.

The similarity between eukaryotic and bacterial STPKs has made the much better understood eukaryotic STPKs a template for understanding STPK signaling in prokaryotes (15,16). In eukaryotes, STPKs engage in often extensive crosstalk that can modulate pathway output, and unlike bacterial TCSs that typically only phosphorylate their specific response regulator, can form branched signaling networks with multiple substrates (17). Bacteria have between none and hundreds of STPKs. For example, *S. cellulosum* codes for 317 STPKs (18), a number close to that found in many eukaryotic organisms that perhaps also implies a similar network organization. However, comprehensive analysis of hundreds of STPKs is challenging, and a systems-wide analysis of a larger bacterial kinome is currently lacking. *Mtb*, on the other hand, expresses eleven STPKs, a number that is small enough to be experimentally tractable yet large enough to potentially give rise to a network of *O*-phosphorylation signaling. A more complex *O*-phosphorylation network structure in *Mtb* was previously suggested by a study of *in vitro* kinase-kinase interactions (19) and initial phosphoproteomic studies using single STPK deletion or chemical inactivation approaches (10,11), but the presence and architecture of a higher-order bacterial STPK *O*-phosphorylation system remains undetermined. Importantly, STPKs are one of the major classes of drug targets in humans, with >60 drugs in the clinic (20). A deeper understanding of *Mtb* STPK function could help translate these successes to this human pathogen that is fueling an enduring global pandemic for which few effective countermeasures are available.

To understand the full ensemble of *Mtb* STPKs and the signaling network they establish, we generated a panel of loss (LoF) and gain-of-function (GoF) STPK mutant strains and comprehensively characterized the STPK’s phosphoproteomic and transcriptional effects. We show that >70% of *Mtb* proteins are *O*-phosphorylated and draw a detailed map of the substrate-kinase relationships of the STPKs, providing >3,700 high confidence substrates of individual STPKs. Remarkably, ∼30% of *Mtb* gene expression was altered by STPK perturbation. This large impact of STPK signaling on transcription could be explained by the pervasive interactions between STPKs, TCSs, and transcription factors (TFs), 54% of which were differentially phosphorylated by STPKs. Together, these data provide the deepest bacterial *O*-phosphoproteome to date, provide thousands of *Mtb* STPK-substrate relationships, identify a large interface between *O*-phosphorylation and transcription in *Mtb*, predict and validate functional STPK-TF interactions, and together establish *O*-phosphorylation as a dominant signaling mechanism with a complexity that is comparable to eukaryotic *O*-phosphorylation systems.

## RESULTS

### An STPK gain-and loss-of-function mutant panel

We sought to comprehensively characterize the *Mtb* phosphoproteome and test the effects of individual STPKs by measuring changes in phosphorylation and gene expression in STPK mutant strains. Nine of the *Mtb* STPKs have an extracellular sensor domain that reaches into the periplasmic space (21). We anticipated that several of these have low basal activity in broth culture and are only fully activated by signals that are likely absent *in vitro*. The loss of such an inactive enzyme would not produce measurable effects. STPKs are typically activated by binding to ligands that promote oligomerization, and this mode of activation appears to be generally conserved in *Mtb* (15) and may be replicated by STPK induction. Thus, we employed a complementary perturbation approach that combines STPK loss-of-function (LoF; knockdown or knockout) and gain-of-function (GoF; induction) to reveal substrates for all STPKs irrespective of their basal activation state.

All mutant strains were generated in the *Mtb* H37Rv background. GoF strains were engineered by inserting a plasmid expressing the complete STPK coding region under control of a tetracycline-inducible promoter. LoF strains were engineered by replacing all STPK genes except for the *in vitro* essential *pknA* and *pknB* by a hygromycin cassette using recombineering (22). Expression of PknB was reduced by CRISPRi (23), with the degree of knockdown titrated so as not to affect cell growth *in vitro*. PknA was not amenable to our complementary strategy, as the PknA kinase activity caused toxicity that precluded expression above normal levels in *Mtb*. Also, a study of the phosphoproteomic consequences of *pknA* deletion was recently presented (11), prompting us to exclude PknA from our study. We grew all cultures to early stationary phase, as slowed replication under nutrient limitation arguably mimics restrictive *in vivo* growth conditions more faithfully than exponential growth, and because the largest degree of *O*-phosphorylation in *E. coli* was reported in stationary phase (24). We confirmed STPK knockout and induction by quantitative, targeted mass spectrometry (MS) using Selected Reaction Monitoring. We observed a range of STPK abundances in WT, with several STPKs present at low concentrations (Fig. 1A). All STPKs were absent or reduced to below the level of detection in the LoF strains, confirming knockout or knockdown, and all GoF mutants showed robust induction (Fig. 1A). In total, we generated 20 STPK mutant strains, a complete LoF/GoF panel except for PknA. To rule out altered viability of the mutant strains that may lead to a non-specific stress or death responses, we tested the viability of mutants in early stationary phase by regrowth after dilution in fresh medium. All mutant strains were viable and grew with wild type (WT) kinetics, except for PknF, which grew slightly slower than WT (Fig. S1).

**Figure 1:**
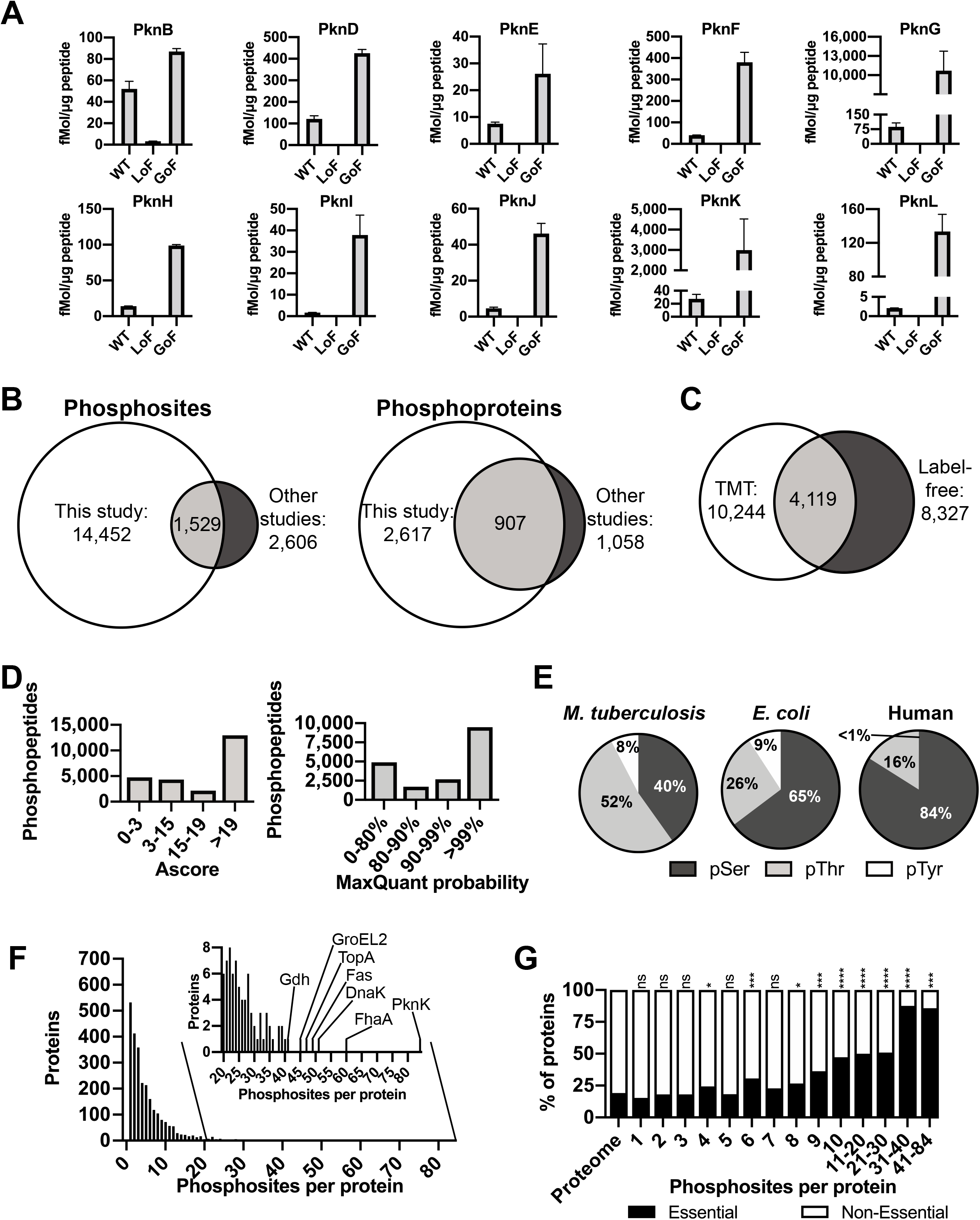
A deep *Mtb* phosphoproteome. (A) Quantitation of STPK protein levels in LoF and GoF *Mtb* mutant strains by selected reaction monitoring. PknB LoF is a knockdown strain, all other LoF strains are knockouts. (B) Comparison of the number of unique protein *O*-phosphorylation sites and -proteins previously published and identified in this study. (C) Number of unique phosphosites identified by TMT and label-free approaches. (D) Distribution of phosphosite localization Ascore (for TMT data) and MaxQuant probability (for label-free data). An Ascore of 19 corresponds to 99% probability of correct localization. (E) Relative share of pSer, pThr, and pTyr sites in *Mtb, E. coli*, and human cells. (F) Number of phosphorylation sites per protein and highly phosphorylated proteins with >20 phosphosites (inset). (G) Share of phosphorylated essential gene products stratified by number of phosphosites. Essentiality was based on transposon mutagenesis (30).

### A deep *Mtb* phosphoproteome

To define a comprehensive *Mtb O*-phosphophoproteome, we analyzed WT and mutant strains by MS-based phosphoproteomics. In pilot studies, we observed that label-free MS identified more total phosphosites, but tandem mass tag (TMT) labeling provided more reliable quantitation, and both methods identified unique phosphosites. To maximize the number of total sites in combination with precise quantitation, we measured all samples by both methods. Together, we analyzed total protein abundance and phosphorylation in three biological replicates for each of the 20 mutant strains and WT by label-free and TMT, >150 phosphoproteomes and >10 million MS^2^ spectra in total. We only considered phosphosites that had a false discovery rate of <0.01% at the peptide level, a PEP score of <0.05, a MaxQuant localization score >0.99 (label-free), or an Ascore of >19 (TMT), corresponding to 99% confidence in phosphosite localization assignment. An overview of the MS workflow is given in Fig. S2.

We detected 3,232 of the ∼4,200 predicted *Mtb* proteins, providing high coverage when compared to previous *Mtb* proteogenomic studies (25) and in light that only ∼80% of *Mtb* genes are expressed at a time (26). We detected 14,452 unique *O*-phosphorylation sites on 2,617 *Mtb* proteins that met our statistical criteria, or 80% of detected *Mtb* proteins. These data show pervasive *O*-phosphorylation comparable to the highest level of phosphorylation detected in eukaryotes, where between 50-70% of proteins are typically found to be phosphorylated (27-29). Our study identified >5 times more phosphosites than all previously published *Mtb* phosphoproteome studies combined (Fig. 1B, Table S1) and represents the largest bacterial phosphoproteome to date. We identified 7,533 pThr, 5,816 pSer, and 1,103 pTyr sites with precise phosphosite assignment. The label-free and TMT approach each added a large number of unique phosphosites, 4,208 and 6,125, respectively (Fig. 1C). The distribution of localization scores for sites detected by label-free and TMT data showed that most sites could be assigned with high confidence, with ∼50% of assignments reaching >99% confidence and over 60% reaching 90% confidence (Fig. 1D). The relative distribution of pSer, pThr, and pTyr detected in our study was 40%, 52%, and 8%, a distinct phosphorylation profile compared to either eukaryotic or the *E. coli* phosphoproteome, where pSer is the most common modification (Fig. 1E). The share of pTyr was higher than that previously detected in *Mtb* (13), and >10-fold higher than in humans (29), indicating a more prominent role of pTyr in *Mtb* (Fig. 1E). Most phosphoproteins (>500) had a single phosphosite, while 413 proteins had two, and 1,330 proteins had between 3 and 10 sites. The number of proteins with more than ten sites was small, but one group of proteins had a remarkably large number of up to 84 phosphorylation sites (Fig. 1F). These highly phosphorylated proteins included PknK, ClpC1, Pks13, and FhaA and could suggest a function of these sites outside of canonical phospho-switches, perhaps based on the physicochemical properties of a large accumulation of negative charges in a cellular subcompartment. We next determined the number of phosphoproteins that are essential as defined by *in vitro* transposon mutagenesis (30). Essential gene products were ∼1.5-fold enriched in the phosphoproteome overall. However, a strong correlation between essentiality and the number of phosphorylation sites was apparent. For example, >36% of proteins with more than nine phosphosites were essential, and ∼88% of those with 31-40 sites were essential (p<0.0001), compared to ∼20% essential genes in the genome overall (Fig 1G).

### The *Mtb* kinase-substrate relationships

We next sought to create a directory of individual STPKs and their cellular substrates. We defined substrates of a given STPK as either hypophosphorylated in that kinase’s LoF or hyperphosphorylated in the GoF mutant strain, or both. Proteins with phosphorylation changes in the opposite direction were considered indirect substrates. To this end, we compared the phosphorylation pattern of each site in every STPK mutant to that of WT and calculated their abundance changes. Due to higher quality quantitation data, we used TMT data for this analysis. We defined a significant change as one detected in all three replicates, changed by at least 2-fold compared to WT, and with a p-value of <0.005. We also included on/off changes with no detected peptides in one set of replicates, although we could not generate p-values for these sites. We identified 3,503 significant changes in unique phosphosites across all mutants, or 24% of all phosphosites (Fig. 2). The overall shifts in the phosphoproteome in response to kinase perturbation were largely as expected, with GoF leading predominantly to increases in phosphorylation and LoF to decreases. Together, we detected 296 and 3,454 unique changes in phosphosites as a result of LoF and GoF, respectively (Fig. 2). The number of substrates varied largely by STPK from 15 to 968 (Fig. 2), suggesting widely varying signaling capacity of the different STPKs.

**Figure 2:**
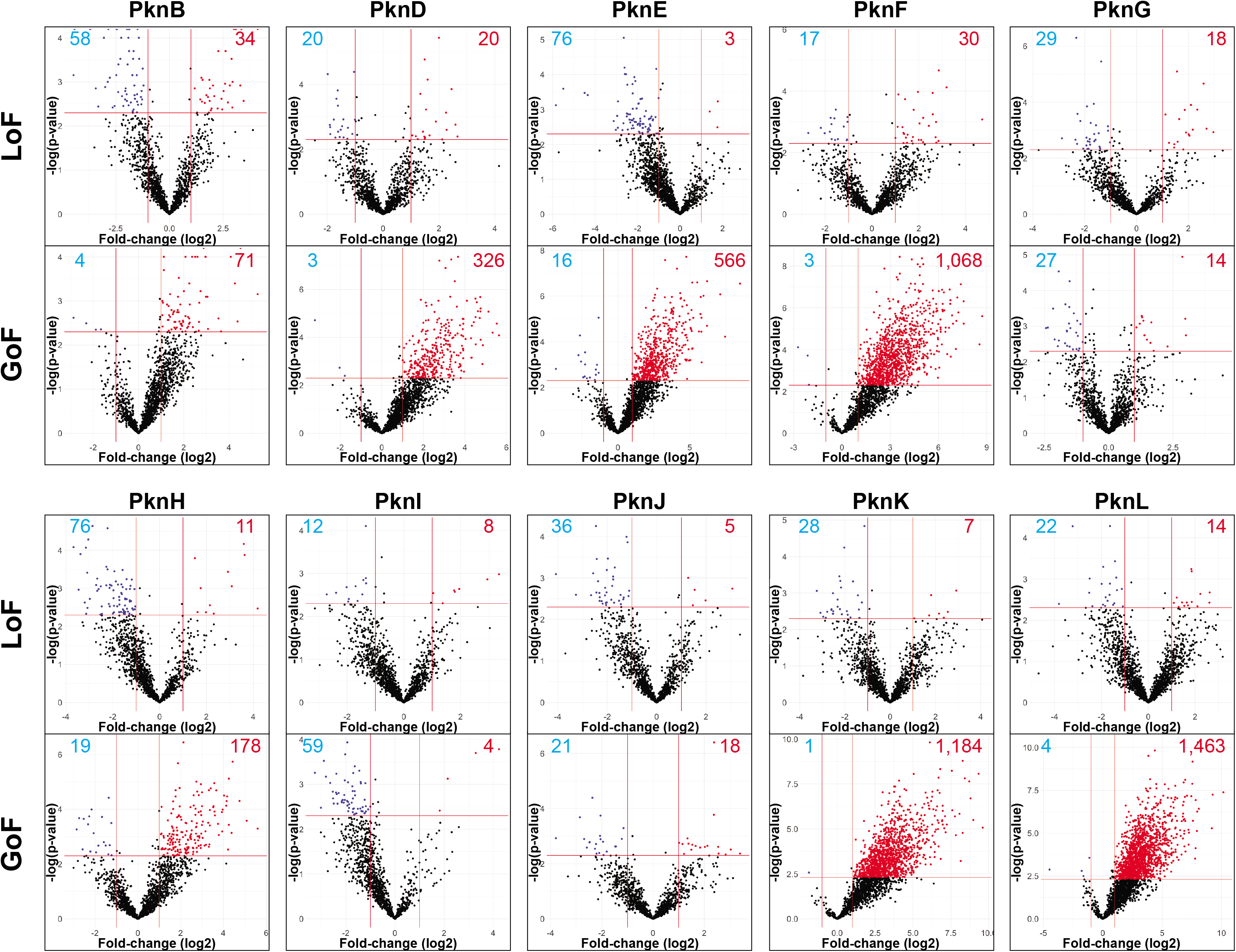
STPK perturbation defines kinase-substrate interactions. Magnitude and direction of changes in phosphorylation sites in LoF (Rows 1 and 3) and GoF strains (Rows 2 and 4). Volcano plots show log_2_-fold changes in phosphorylation levels by phosphosite (X-axis) versus the −log p value of the change (y-axis). Horizontal and vertical lines show significance cutoffs applied to the data for further analysis (>2-fold change in abundance, p<0.005). Significantly changing phosphosites are shown in red and blue and their numbers are given in the respective quadrant. The STPKs altered in the respective strain were not included as they showed the largest changes and distorted the plots.

Consistent with our premise, many STPKs had low basal activity in the WT strain in broth culture. PknD, F, I, K, and L showed little substrate hypophosphorylation (<30 phosphosites) when deleted (Fig. 2, 3A). However, these silent kinases could be effectively activated to phosphorylate substrates in the GoF strains (Fig. 2, 3A), allowing for the definition of a distinct substrate set for each STPK (Table S2). We next tested if STPK induction in the GoF strains may lead to off-target phosphorylation. We found that the average magnitude and number of phosphorylation changes did not correlate with the abundance of the respective kinase (Fig. S3A). For example, the PknG GoF strain, which showed the highest degree of STPK induction, generated only 41 phosphorylation changes. These data are consistent with GoF leading to predominantly physiologic phosphorylation events. With a total number of >14,000 unique phosphosites, each of the STPKs would on average be expected to phosphorylate ∼1,400 substrates. Only the PknK and PknL GoF strains phosphorylated as many substrates, indicating that induction in our strains may still not fully capture the full activity for most STPKs.

**Figure 3:**
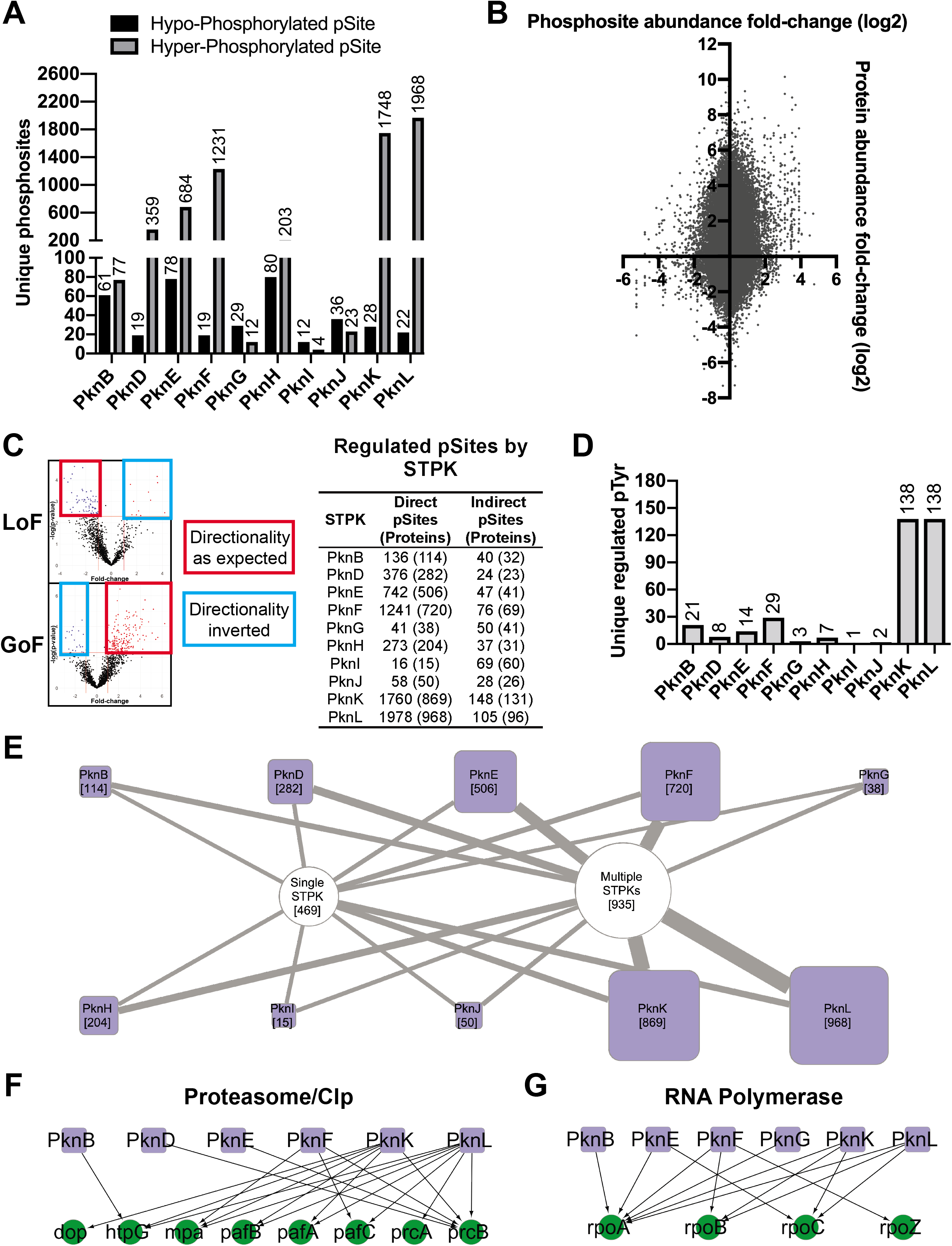
*Mtb* protein *O*-phosphorylation establishes a network. (A) Effects of STPK perturbation on the abundance of unique phosphosites by STPK compared to WT. Changes in LoF and GoF strains were combined for each STPK. (B) Correlation between global protein abundance and phosphosite abundance of all phosphorylation sites showing a change in STPK LoF and GoF mutants. Quantitative TMT data of total protein abundance was plotted against the phosphosite abundance for each protein, showing that most increases in phosphorylation are not concomitant with higher protein abundance. (C) Directionality of phosphorylation changes in response to STPK perturbation. The volcano plots show assignment of direct (hypophosphorylation in LoF, hyperphosphorylation in GoF), and indirect phosphorylation events with inverted directionality. Table shows putative direct and indirect phosphosites (phosphoproteins) by STPK. (D) Changes in Tyr phosphorylation sites upon STPK perturbation. The numbers given above the STPK mutant strain is from LoF and GoF mutants combined. (E) An overview of the STPKs’ substrates and overlap in substrate phosphorylation. The purple squares represent the STPKs. The size of the square is proportional to the number of that STPK’s substrates, which is also given in parentheses. The round nodes show the number of substrates phosphorylated by one or multiple STPKs. The thickness of the edge corresponds to the number of substrates phosphorylated by the respective STPK. (F) Phosphorylation of members of the proteasome/Clp protein degradation systems by different STPKs. (G) Phosphorylation of RNA polymerase subunits by multiple STPKs.

Differential phosphorylation can result from changes in phosphoprotein abundance and/or from changes in occupancy of phosphosites. To understand the relationship between protein abundance and phosphosite occupancy, we determined the total abundance of proteins in all strains by MS and correlated abundance with phosphorylation (Fig. 3B). There was no significant correlation between the two (r=0.09) indicating that changes in phosphorylation primarily occur on the level of site occupancy. Kinases interact with each other more than with other protein families and can profoundly affect each other’s activity by phosphorylation (17,19). A corollary of these interactions is that phosphorylation events detected upon kinase perturbation may be indirect, which is difficult to discriminate from direct effects (31). To identify indirect phosphorylation events, we examined the number of hyperphosphorylated sites in the LoF mutants and of hypophosphorylated sites in the GoF mutants as these cannot be explained by a direct STPK-substrate interaction (Fig. 3C). While all STPKs showed some of these indirect effects, their number was ∼8% of all altered phosphosites (Fig. 3C). All indirect sites are indicated in Table S2. Additional indirect substrates might also be among the substrates hypophosphorylated in the LoF and hyperphosphorylated in the GoF mutant, but these are indistinguishable from direct substrates in our data.

To identify potential substrate sequence preferences of the STPKs, we compiled the substrates that showed significant changes in LoF and/or GoF mutants and identified the amino acids most commonly found adjacent to the phosphorylation site using the pLogo tool (32) (Fig. S3B). The *Mtb* STPKs’ preferences were different from the often basophilic and proline-directed motifs in eukaryotes and were generally less well-defined (33,34). Motif preferences were largely shared among the pSer and pThr substrates, but not the pTyr substrates. Motifs associated with pSer and pThr sites showed preferences in the −1 (D), +1 (G/D/E), +2 (P/E) position relative to the phosphosite (Fig S3B). Charged residues (D, E, K) were generally favored in the −5 to +5 positions, and hydrophobic residues were disfavored in most positions. The preferred pTyr motif was less clearly defined, with moderate preference for G/Q/D in −1 and hydrophobic residues (V, L, I) in the +1 positions. To further parse individual STPKs’ motifs, we mapped preferred sites of the STPKs individually. Motifs of many STPKs predictably reflected the overall preference, and motifs were generally weak, suggesting other substrate selectivity determinants than the primary sequence of the phosphosite (data not shown).

### Several STPKs are dual specificity kinases

Protein Tyr phosphorylation was long thought to be absent in *Mtb* but has now conclusively been shown (13). Given that canonical bacterial Tyr kinases are absent in *Mtb*, the identity of the Tyr kinases remained an open question. Based on *in vitro* biochemical activity, an atypical kinase and dual specificity kinases among the STPKs have previously been suggested as candidates (13,35). Our data further expanded the number of pTyr sites in *Mtb* and provided the opportunity to test the hypothesis of dual specificity of the STPKs in a cellular context. While all STPKs showed a preference for phosphorylating Ser/Thr over Tyr residues, many also phosphorylated Tyr residues. The pTyr sites affected by STPK permutation account for 231 pTyr sites, or 24% of all pTyr sites detected in our study. In particular PknK and PknL LoF and GoF mutants showed a marked decrease and increase of pTyr, respectively (Fig. 3D). These data suggest that some *Mtb* STPKs have evolved dual activity similar to the dual specificity kinases in many eukaryotes to perform a protein modification that is typically executed by specialized bacterial Tyr kinases.

### The STPKs constitute a distributed network

We next analyzed the STPK-substrate relationships globally to detect larger features of the network by visualizing all STPK-substrate interactions. While 469 substrate proteins were only phosphorylated by a single STPK, 935 were phosphorylated by more than one STPK (Fig. 3E). Of the proteins phosphorylated by more than one STPK, most were phosphorylated by two STPKs (292), and one, Rv0020c, was phosphorylated by all ten STPKs. The high redundancy of this phosphorylation network may explain the relative paucity of *in vitro* phenotypes of KO strains of most of the STPKs despite their large number of substrates and is reminiscent of distributed eukaryotic phosphosignaling networks (36). We next inspected the STPK substrates showing significant changes upon kinase perturbation for enrichment in several protein families and GO terms. Among the most significantly enriched were components of the proteasome and Clp protease systems (Fig. 3F), and RNA Polymerase (Fig. 3G), almost all of which were phosphorylated by multiple STPKs suggesting coordinated regulation by several STPK inputs of these central cellular functions.

### STPKs have global transcriptional effects

We identified 158 phosphorylated TFs (77% of *Mtb* TFs (37)) and 112 TFs (54% of TFs) with significant phosphorylation changes associated with STPK perturbation, suggesting potentially widespread regulation of TF activity by *O*-phosphorylation (Fig. 4A). In a simple transcriptional relay, an STPK and gene expression can be directly linked by the phosphorylation of a TF, leading to gain or loss of DNA binding (38). The extensive phosphorylation of TFs predicted that the transcriptional effects of the STPKs could be significant. To test this idea, we measured the STPKs’ downstream transcriptional effects by RNA-seq using samples matched to the samples used for phosphoproteomics. We used a cutoff of at least 4-fold up-or down-regulation compared to WT and a significance cutoff of p <0.01 to define differentially expressed genes (DEGs).

**Figure 4:**
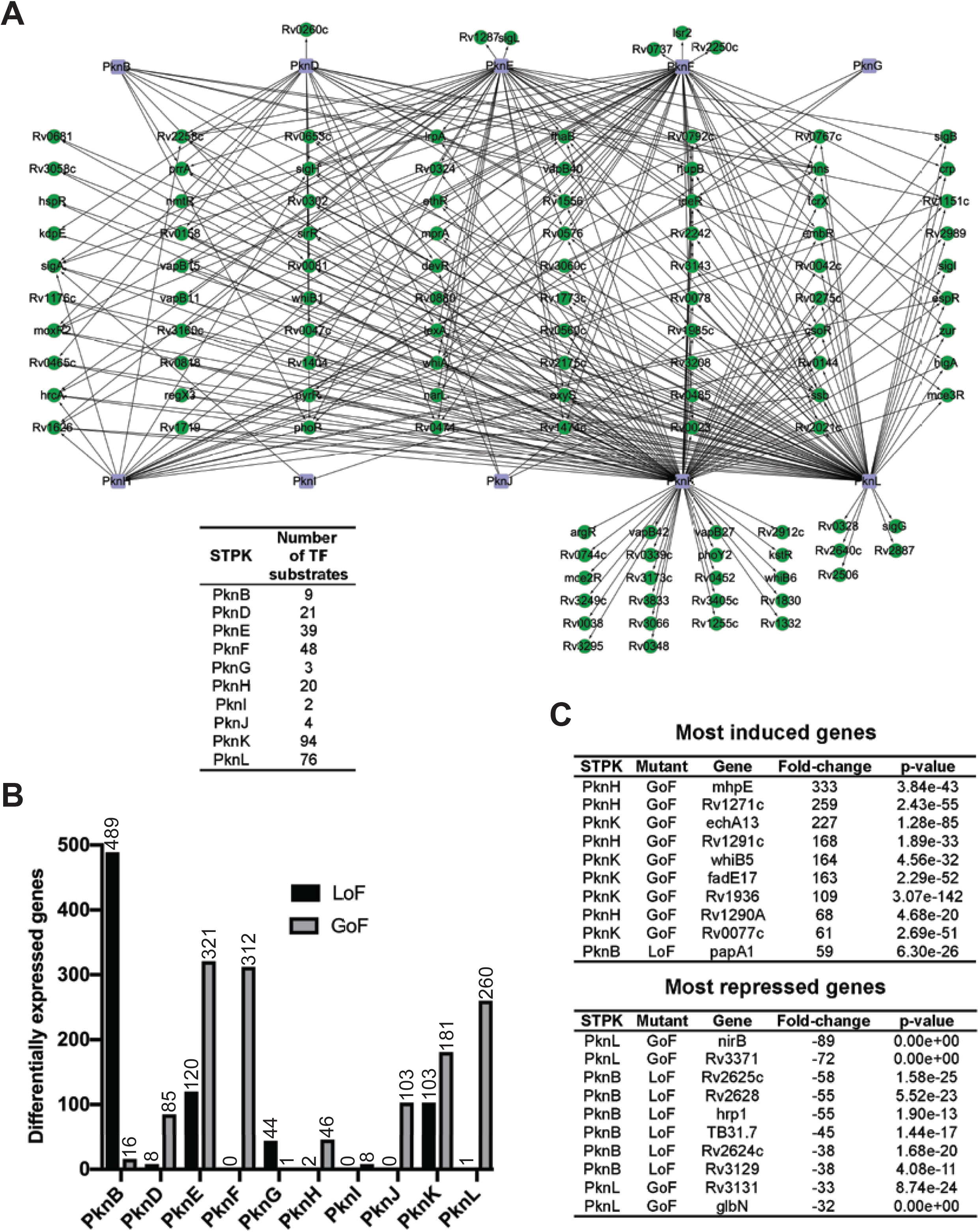
*O*-phosphorylation has large transcriptional effects. (A) Overview of TF phosphorylation by STPKs. STPKs are shown as purple squares, TFs are shown as green circles. Edges represent kinase-substrate interactions. No Ascore cutoff was applied as analysis was on the basis of phosphoproteins, not phosphosites. Table shows number of TF substrates for each STPK. (B) Overall number of differentially expressed genes upon STPK perturbation by STPK and by type of perturbation. Differentially expressed genes are defined as >4-fold change with p<0.01. (C) The most induced and repressed genes by STPK and type of perturbation.

Overall, 1,155 of 4,030 detected genes met our criteria for DEG, representing ∼30% of *Mtb* genes (Fig. 4B, S4). All STPKs caused changes in gene expression and two distinct kinase activation patterns were apparent: Several STPKs showed a larger change in gene expression upon LoF (PknB, G), suggesting a constitutive housekeeping function. In particular PknB elicited differential expression of 30-fold more genes in the LoF strain than the GoF strain (Fig. 4B, S4). Another group of STPKs including PknD, F, H and J showed no or little expression changes upon LoF, but large changes upon GoF (Fig 4B, S4). These data mirror findings from our phosphoproteomic analysis showing that some STPKs are inactive in culture and likely require a signal for activation that is absent *in vitro*. Except for PknE and PknF, the number of differentially regulated phosphosites and the number of DEGs corresponded well (r=0.92) (Fig. S5A). The range of fold-change varied widely to several hundred-fold (Fig. 4C). Although proteomic analyses described above suggested no or little artificial phosphorylation of the GoF mutants, we re-examined this question using transcriptional data. To test if the abundance of STPKs in the GoF mutants was within physiologic levels, we mined existing transcriptional data for STPK expression during stress conditions and found that in most cases, STPK expression levels in the GoF mutants were within the range found in WT bacteria subjected to physiologic stresses such as hypoxia, oxidative stress, and nitrosative stress (Fig. S5B). The PknK GoF mutant showed the largest difference from PknK induction under stress in WT, however, the levels of induction of 8 STPKs in our GoF strains were in fact below those reported in WT under specific stresses. Non-specific transcriptional effects of the STPKs would also suggest a correlation between the magnitude of STPK expression and the number of DEGs. The number of differentially expressed genes was not correlated with induction of the STPK (r=0.37) (Fig. S5C). To test if the transcriptional effects of STPK perturbation are indeed due to the kinase activity of the STPKs, we measured the transcriptional effects of a kinase-dead PknL mutant (Fig. S6A). While the PknL GoF mutant led to 260 changes in gene expression compared to WT, the kinase dead mutant led to one, showing that the kinase activity is responsible for gene expression changes. Domain prediction showed that in addition to the kinase domain, PknK also has a transcriptional regulator domain of the MalT family. To test if gene expression changes in the PknK GoF strain data reflect effects of the kinase activity or perhaps also effects of the MalT domain, we compared DEGs in strains overexpressing PknK WT and kinase dead mutant. The WT GoF mutant showed changes in 181 genes, the kinase dead mutant in 13, suggesting that <10% of DEGs in the PknK GoF mutant may be due to the activity of the MalT domain (Fig. S6B).

To understand the higher-level organization of the STPKs’ transcriptional effects, we visualized the DEGs regulated by STPK LoF and GoF mutants (Fig. S7). While some genes were affected by a single STPK, 40% of genes were affected by several STPKs, some by up to seven. The co-regulation of these genes showed that signaling by multiple *Mtb* STPKs converges on the same cellular pathways and suggests a high degree of redundancy between the STPKs. We observed co-regulation in the same and in the opposite direction. For example, Rv2590 was induced in PknG and PknL LoF mutants while Rv1875 was repressed in PknE and PknK LoF mutants, both cases of redundant kinase function. In contrast, Rv1739c was upregulated in the PknE LoF mutant and downregulated in the PknB LoF mutant. The connectedness of STPK gene expression given as the average number of STPKs that affect expression of one gene was on average 1.75. These data reveal a large, highly coordinated interface between STPK signaling and transcription.

### Integration of phosphoproteomic and transcriptional data predicts functional TF phosphosites

The share of functional phosphorylation events in any phosphoproteome is unknown, and their prediction remains a major challenge. Because we determined STPK-TF substrate pairs and global functional readouts for the STPKs by measuring transcriptional changes, our data offered an opportunity to predict function of phosphorylation events on TFs. To this end, we integrated our phosphoproteomic data on STPK-TF kinase-substrate pairs, the matched STPK transcriptional data, and a global transcriptional dataset mapping the transcriptional effects of ∼200 *Mtb* TFs (37). A significant overlap between the regulon of an STPK and that of its cognate TF substrate would suggest that the phosphorylation site is functional. To correlate regulons of phosphorylated TFs and those of their cognate STPKs, we determined the genes altered by an STPK and its TF partner and plotted the % overlap of the TFs regulon with the STPK’s regulon (Fig. 5). 52 TFs showed significant regulon overlap (p=0.05) with their cognate STPK’s regulon. 24 TFs showed >50% of shared regulated genes, and the regulons of 3 TFs (hns, pyrR and Rv0792c) were fully contained in their STPK’s regulon. These data predict a set of TFs whose phosphorylation is likely regulatory and leads to downstream gene expression changes. The strength of the predictions increase with higher regulon overlap and lower STPK regulon size (Fig. 5).

**Figure 5:**
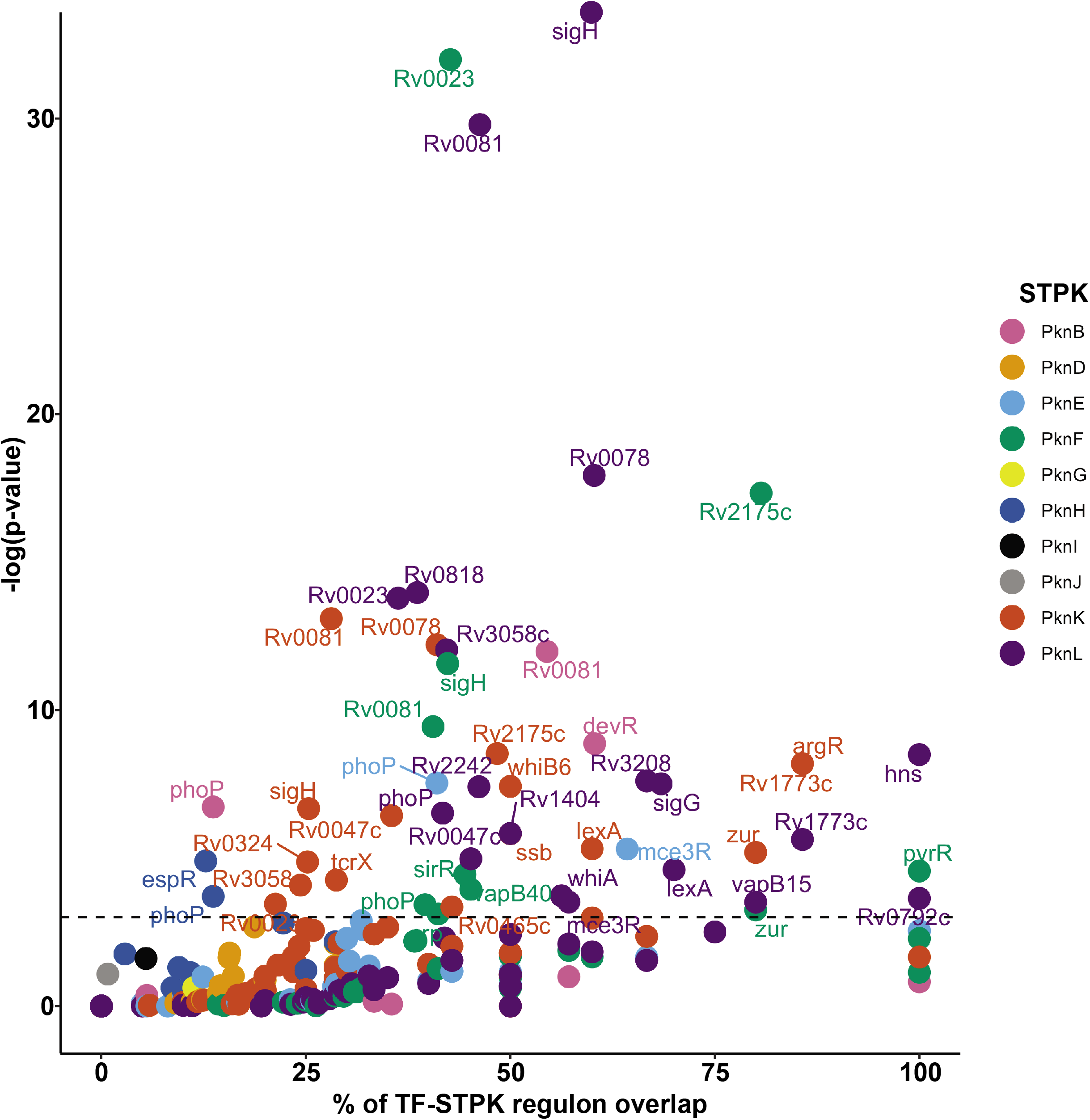
Prediction of functional TF phosphorylation events. Genes regulated by both TFs and their cognate STPK were plotted as % regulon overlap against the significance of the overlap. Colors denote different STPKs. Dotted line indicates a p-value of 0.05. These data are based on protein-level phosphorylation and no localization score cutoff was applied.

### *O*-phosphorylation alters Zur transcription factor activity

To validate the STPK-TF functional predictions and to illustrate how these predictions can be used to assemble signaling pathways, we explored the interaction between PknK and the Zinc uptake repressor Zur (Rv2359), a negative transcriptional regulator of metal homeostasis (39).

The PknK GoF mutant showed significant Zur hyperphosphorylation on Thr67 in our MS data (8.7-fold, p-value=0.001) when compared to WT, consistent with a direct kinase-substrate interaction, and the phosphorylation event was predicted to be functional based on significant regulon overlap in our STPK-TF regulon analysis (Fig. 6). We validated the PknK-Zur interaction using an *in vitro* kinase assay with recombinant PknK and Zur. PknK efficiently phosphorylated Zur *in vitro* (Fig. 6B), but PknG, which did not affect Zur phosphorylation in our phosphoproteomic data, did not. To test a regulatory connection between PknK and Zur, we first compared the previously identified Zur LoF and GoF regulons (37,39) to genes regulated in the PknK GoF mutant and found that >80% of the Zur regulon is also regulated in the PknK GoF mutant, suggesting a function in the same pathway (Fig. 6C). While the Zur GoF mutant strain repressed its regulon consistent with its role as a negative regulator, the PknK GoF mutant strain led to induction of nearly all genes in the Zur regulon. The opposite effects of Zur and PknK GoF predicted that phosphorylation of Zur by PknK derepresses Zur. Structural analysis using a high-confidence model based on the *E. coli* Zur structure bound to DNA showed that Thr67 is adjacent to the DNA-binding helix-loop-helix motif of Zur and faces directly towards the minor groove of target DNA. In its phosphorylated form, Thr67 would likely create charge repulsion with the phosphatedeoxyribose backbone of the DNA, predicting a negative effect of phosphorylation on DNA binding. To test the effect of Zur Thr67 phosphorylation on its ability to bind DNA, we purified recombinant WT, a Thr67 phosphoablative (Thr67Ala) and phosphomimetic (Thr67Asp) mutant and tested DNA-binding activity by electrophoretic mobility shift assays (Fig. 6D). WT Zur efficiently bound to DNA, resulting in a shift of the DNA probe.

**Figure 6:**
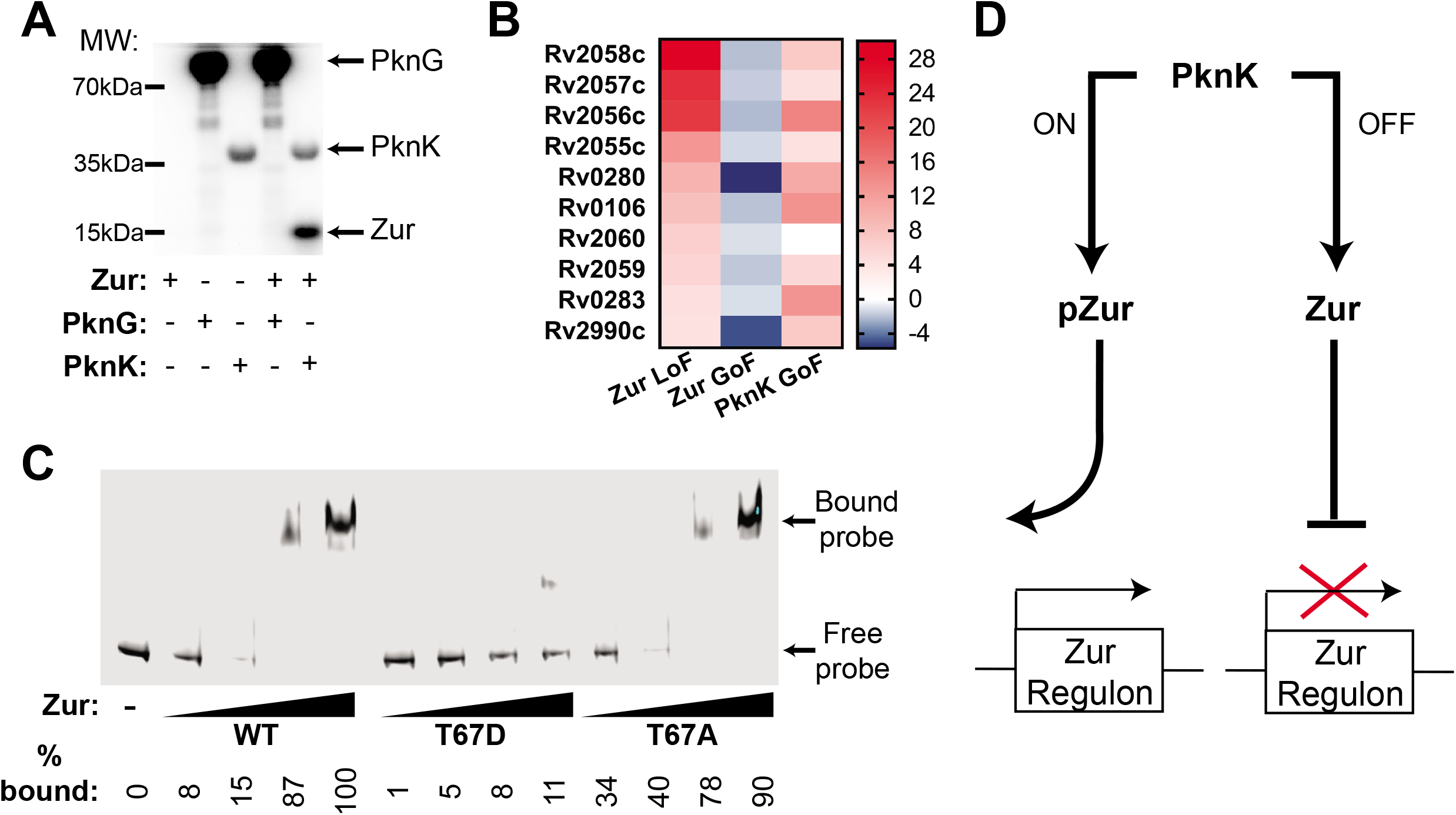
PknK regulates the transcription factor Zur. (A) *In vitro* phosphorylation assay showing specific phosphorylation of recombinant Zur by PknK, but not PknG. (B) Transcriptional response of Zur LoF, Zur GoF, and PknK GoF mutations on the Zur operon shows overlapping but opposite effects of PknK and Zur GoF, predicting derepression of Zur activity by PknK phosphorylation. (C) Effect of Zur Thr67 phosphoablative and -mimetic mutations on DNA binding activity. (D) Model of the PknK-Zur pathway.

The phosphoablative mutant did not show any difference to WT in DNA-binding. The phosphomimetic mutant, however, showed decreased DNA-binding, consistent with our prediction. Together, these data define a phosphoregulation pathway in which PknK phosphorylates Thr67 of Zur, resulting in loss of DNA binding, which de-represses the Zur regulon (Fig. 6E).

### A combined transcriptional and phosphoproteomic resource for studying kinase function

The matched kinome-wide phosphoproteomic and transcriptomic datasets provide the set of phosphorylation sites and genes that are affected by individual STPKs. An overview of the combined interactions is shown in Fig. 7. Our data assign STPKs to substrates and genes and predict STPK functions. Conversely, these data allow for the identification of candidate STPKs affecting a particular physiologic response if that response is characterized by the expression of a gene set or a change in phosphorylation. In addition to several STPKs for which no experimental functional data is available to date, in particular the GoF data furnishes a large set of phosphosites that were not accessible before. The complete dataset showing all transcriptional and phosphorylation effects of the STPK GoF and LoF mutants are presented in Table S1 (all phosphosites, label-free and TMT), Table S2 (regulated phosphosites), and Table S3 (transcriptional changes). Together, these data show that a large swath of *Mtb* physiology is integrated with STPK signaling, provide a resource for assigning kinases to substrates and regulated genes and thus the foundation for detailed mechanistic understanding of this pervasive *Mtb* phosphosignaling mechanism.

**Figure 7:**
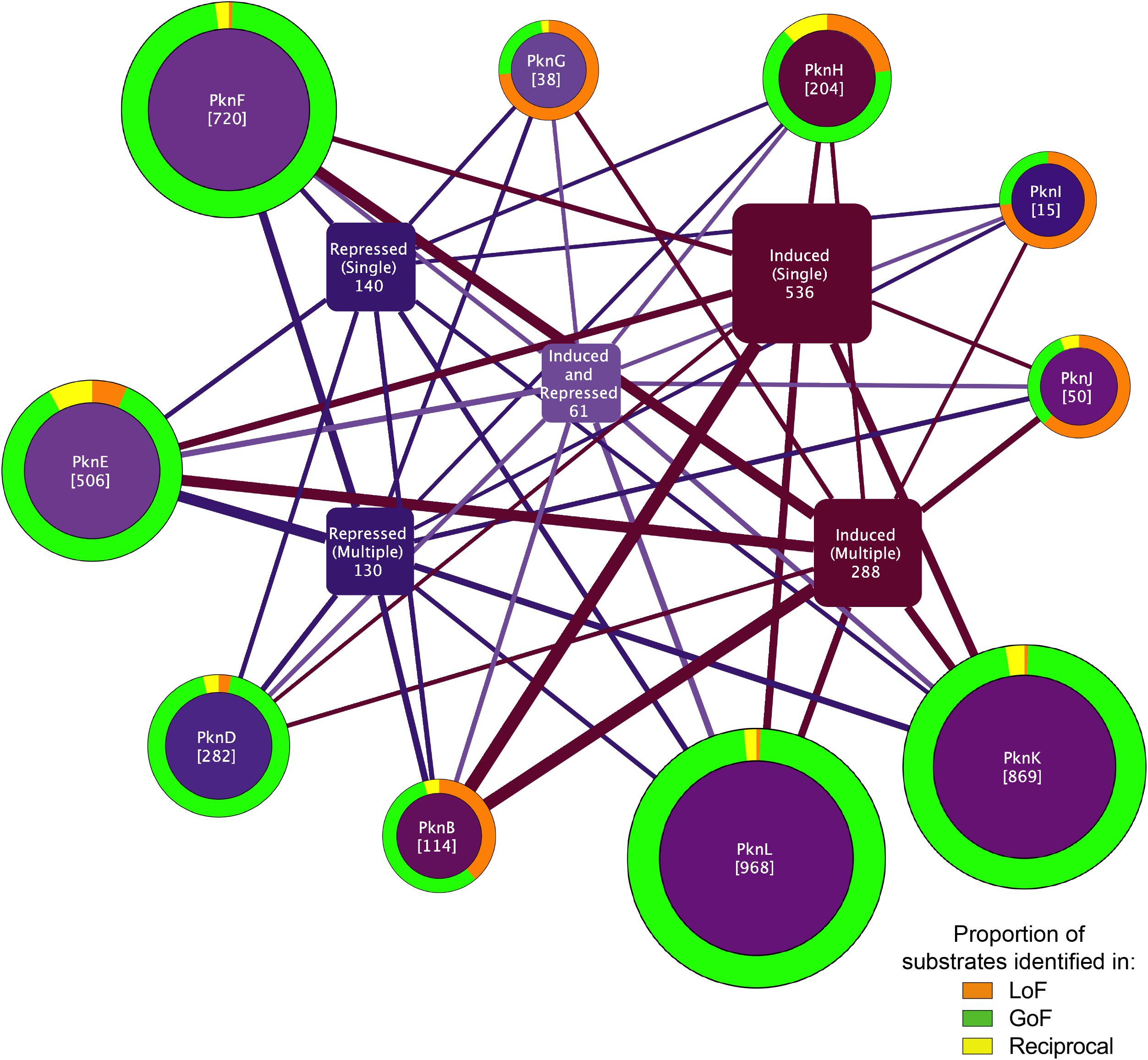
Overview of the *Mtb* STPK *O*-phosphoproteome and transcriptional network. The phosphoproteomic and transcriptional effects of STPK mutation are shown, illustrating the strongly overlapping nature of the network. The STPKs are shown as round nodes, with their total number of phosphorylated substrates in parentheses. The outer circle of each STPK node shows the relative share of substrates observed in the LoF (orange), GoF (green), or both (down in LoF, up in GoF, yellow). The color of the inner STPK circle indicates whether the STPK’s effect on DEGs on average is induction (red), repression (blue), or a combination (purple). The inner square nodes represent genes regulated by the STPKs, with the thickness of edges connecting the STPKs to genes corresponding to the number of genes regulated by an STPK. Single and multiple refers to the number of STPKs that have the particular effect on the group of genes (for example, 130 genes are repressed by more than one STPK). Interestingly, the expression of 61 genes is activated by some STPKs and repressed by others.

## DISCUSSION

The contribution of protein *O*-phosphorylation to bacterial cell signaling remains an open question, and the relatively small numbers of *O*-phosphorylation sites previously reported in bacteria have reinforced the notion that bacterial *O*-phosphorylation systems are simpler and more rudimentary than its eukaryotic counterparts. Here, we challenge this view with the deepest phosphoproteome measured in a bacterium yet, which expands the number of reported *Mtb O*-phosphorylation sites by >5-fold. The degree of *O*-phosphorylation in *Mtb*, with >70% of proteins decorated by at least one phosphosite, is on par with that found in many eukaryotic organisms (28), and shows a signaling network of a complexity that was thought to be exclusive to eukaryotes. These findings place *O*-phosphorylation at the center of cellular signaling in the pandemic human pathogen *Mtb* and argue that the STPKs constitute the backbone of bacterial phosphosignaling in *Mtb*.

To further understand phosphosignaling beyond the general inventory of the phosphoproteome requires parsing of contributions of individual kinases and their substrates, or the kinase-substrate relationships. Assigning these kinase-substrate relationships remains a major experimental bottleneck. Although several substrates, partial pathways, and functions of the *Mtb* STPKs have previously been identified, they remain unknown for most. Global phosphoproteomic approaches are arguably best suited to identify kinase-substrate relationships. These global approaches typically use LoF perturbations and then measure the resulting changes in the phosphoproteome by MS (40). While the changes resulting from a LoF perturbation are highly informative, this approach requires a baseline activity of a kinase in the experimental condition, which was an untested premise for most *Mtb* kinases grown in broth culture. Given that *Mtb* STPKs are mostly sensor kinases that likely survey the host, it is not surprising that many are largely inactive in broth culture, where host-specific signals are absent. Our GoF approach captured many of these previously intractable phosphorylation events and downstream transcriptional effects of these silent STPKs.

Our study vastly expands the overall number of *Mtb O*-phosphorylation sites and candidate substrates for the individual STPKs, including for PknB that has been the subject of dozens of studies including one that applied LoF-MS-based approaches that are comparable to ours (11). The higher number of phosphosites identified in our study is likely a result of high-sensitivity MS hardware, using label-free and TMT approaches combined with extensive sample fractionation, and the analysis of GoF mutants. The extensive substrate sets for the STPKs presented here provide a directory for identifying STPK pathways and functions. One global functional consequence of STPK signaling in particular was the sweeping effect on gene expression, with ∼30% of all *Mtb* genes affected by one or more STPKs. The most direct connection between STPKs and gene expression is through the phosphorylation of TFs, whose DNA binding activity can change upon phosphorylation. Here, we uncover extensive phosphorylation of TFs by the STPKs. A key question for these and in fact most known phosphosites is whether they are functional. Prediction of functional phosphorylation sites is exceedingly difficult. Our data, by measuring both STPK-TF pairs and transcriptional effects, allowed for the prediction of functional phosphorylation sites on TFs and identifies putative signaling pathways from kinase to gene. Most TF phosphorylation events, however, did not produce significant gene regulatory changes that could be ascribed to its cognate STPK. The lack of detectable function of these TF sites may be due to the limited, one-directional perturbation in the TF dataset (37), indirect effects of STPK and/or TF, functions other than transcription, but likely also reflects a large degree of phosphorylation with no function. These data provide a rare experimental datapoint for gauging the share of functional phosphorylation sites within a large protein family.

The many STPK-TF connections predicted to be functional and the regulation of Zur by PknK suggest a general model of STPK-mediated gene regulation in *Mtb* —direct regulation of TFs by STPKs. The number of these interactions was surprising given the few previous examples of STPK-TF interactions reported in *Mtb*. Other direct interactions between STPKs and the transcriptional machinery suggested by our data include the STPK-specific phosphorylation of sigma factors, RNA polymerase, but also of TCSs. TCSs and STPKs were thought to be largely insulated from one another, but the extensive phosphorylation of His kinases and response regulators by the STPKs suggest close integration of the two arms of phosphosignaling. The complex and highly redundant architecture of the STPK signaling network may be better suited to translate complex signals into effective and robust responses than the typically simpler and more linear TCS pathways. The remarkable interconnectivity between signal transduction and gene regulation underscores how understanding the interplay of these networks will be critical to dissect how *Mtb* and potentially other bacteria sense, process, and respond to environmental cues.

MS data are probabilistic in nature. We sought to minimize the uncertainty inherent in our phosphoproteomic data by setting stringent cutoffs for peptide identification, false discovery rate, and phosphosite localization. One limitation of our study is the analysis of the phosphoproteome in only one *in vitro* condition, stationary phase. In addition, gene KO can lead to compensatory effects that can obscure STPK functions. Further, the large effects of the STPK GoF mutants on phosphorylation and transcription also raise the question of experimental artifacts. However, we found no evidence for capricious phosphorylation. The induction levels of STPKs in the GoF mutants were within the range elicited in WT responses to physiologic stress conditions, and the frequency of phosphorylation did not correlate with STPK expression levels. In addition, most changes in the GoF strains were in fact changes in the abundance of phosphosites that were also present in WT. The likelihood of an inadvertent functional impact of erratic phosphosites appear small: Phosphosignals propagate through evolved protein-protein interactions that are unlikely to be replicated by artificial phosphorylation events, and the low occupancy of erratic phosphorylation would further reduce phenotypic consequences. However, a phosphosite or phenotypic effect that can be attributed to a LoF mutation may ultimately be more reliable than an effect observed in a GoF mutant, and corresponding effects of LoF and GoF mutants can further increase the confidence in the detected change.

On a systems level, this study shows a central and dominant place of *O*-phosphorylation in the important human pathogen *Mtb* and a cooperative signaling system of a complexity that is typically only associated with eukaryotes. These findings revise, at least for *Mtb*, the long-held view of the TCSs as the canonical bacterial phosphosignaling system. On a molecular level, this study provides rich detail of this system through a resource with >14,000 unique phosphosites and >3,700 STPK-specific phosphosite changes and their transcriptional effects. These data lay a foundation for understanding the cellular functions of these enzymes and provide signaling context for a large part of *Mtb* physiology.

## SUPPLEMENTAL FIGURE AND TABLE LEGENDS

**Figure S1:**
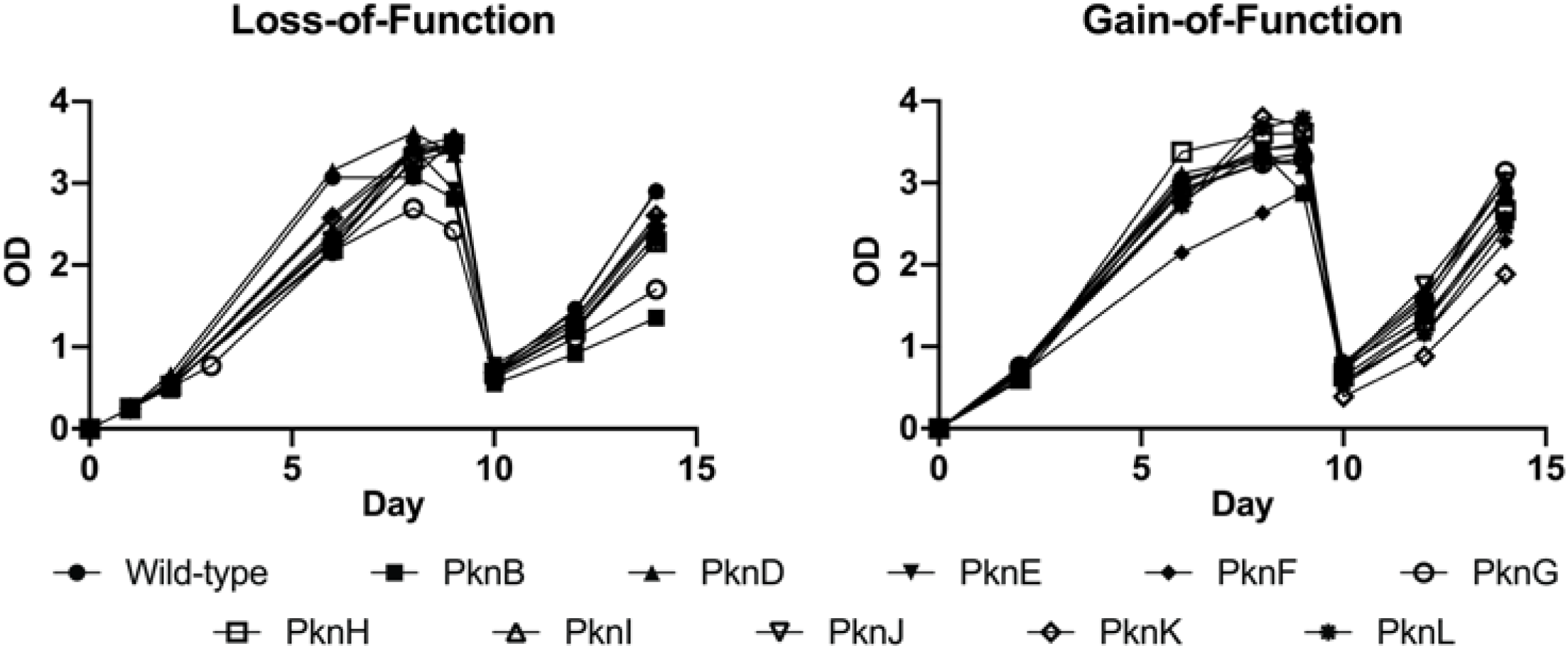
Overview of the MS experimental workflow.

**Figure S2:**
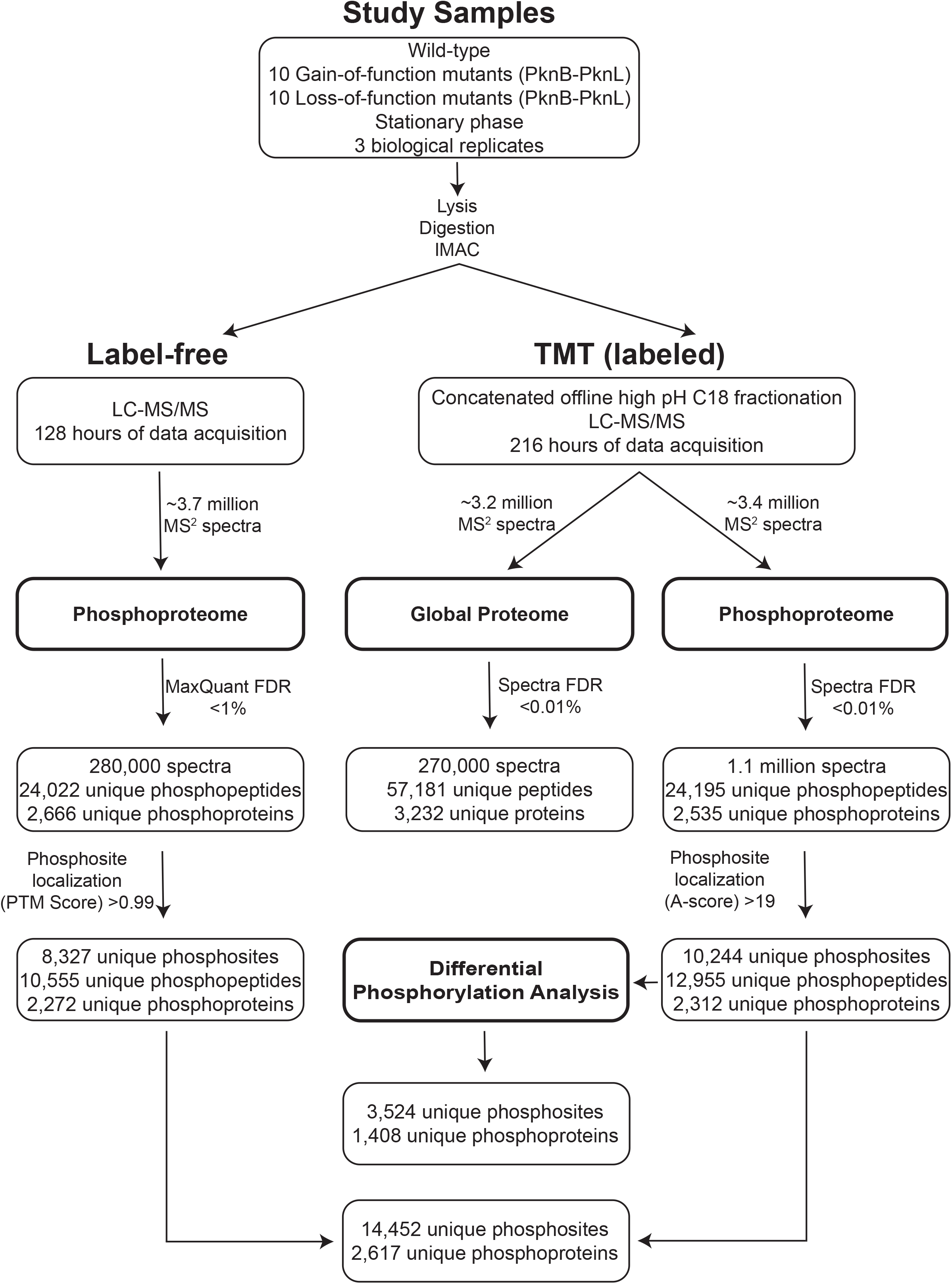
Fitness test of STPK mutant strains. (A) LoF strains were grown to stationary phase, diluted to OD_600_ of 0.25 and regrown to test for fitness. The PknB LoF knockdown mutant was induced with ATc. (B) GoF mutants were induced with ATc, grown to stationary phase, diluted to OD_600_ of 0.25, and regrown without ATc.

**Figure S3:**
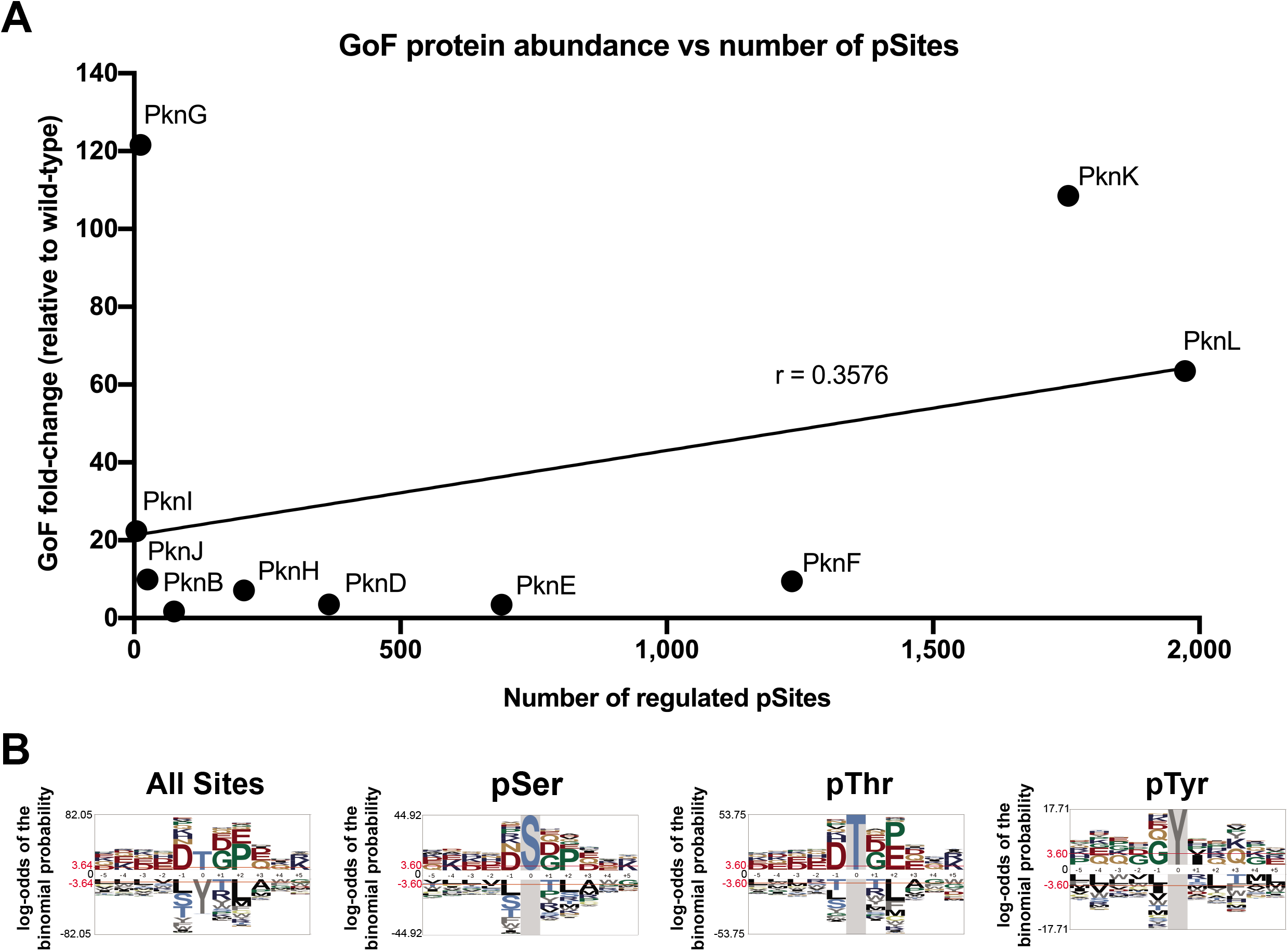
Correlation of phosphosites and STPK protein abundance and motif analysis. (A) The abundance of STPKs in the GoF mutants does not overall correlate with the number of phosphosites detected in these strains. (B) Preferred phosphorylation motifs of all STPKs combined and by phosphoresidue. Height of sequence logo represents frequency of residue at that position.

**Figure S4:**
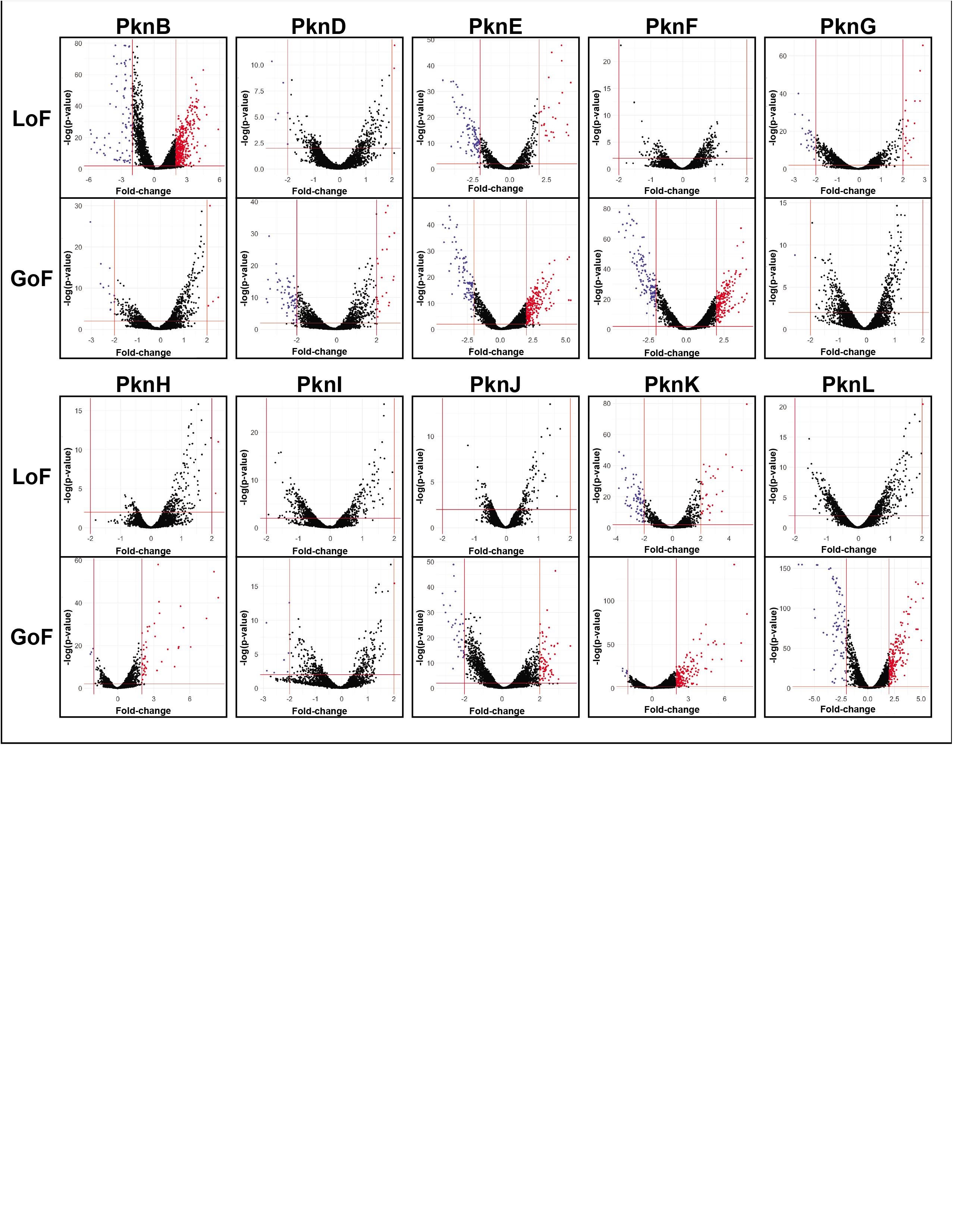
Transcriptional effects of STPK perturbation. Volcano plots show the DEGs in STPK mutant strains, direction of change, magnitude, and p-value associated with the change. Horizontal and vertical red lines show significance cutoffs (>4-fold, p<0.01) used for further analysis of DEGs. The STPKs that were altered in the respective strain were removed from the data as they showed the largest changes and distorted the plots.

**Figure S5:**
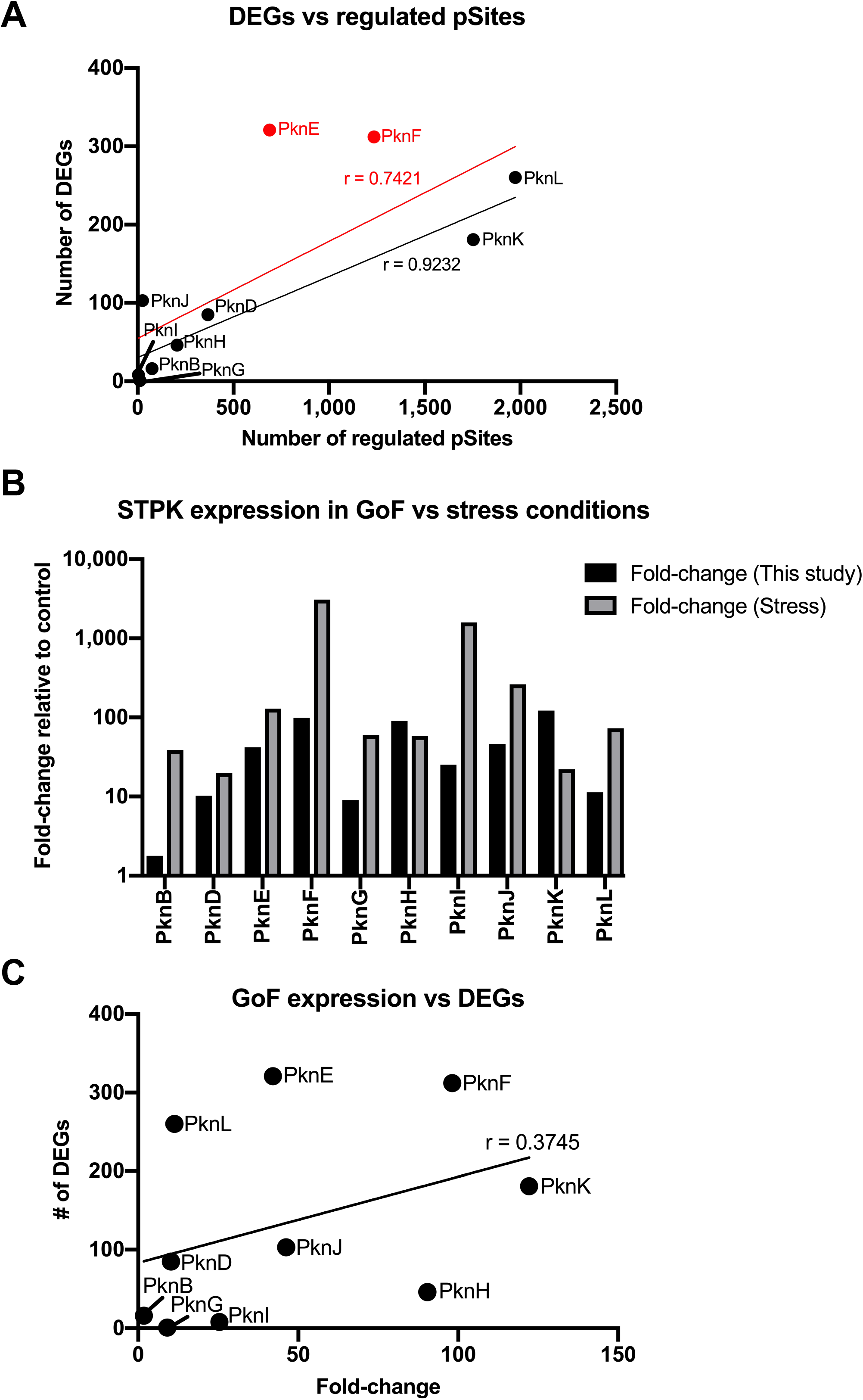
Relationship between STPK induction, phosphosites, and DEGs. (A) More phosphorylation sites in the GoF mutant strains led to more DEGs. PknJ affects the expression of a large number of genes through few phosphosites. Red line shows linear regression if outliers PknE and PknF are excluded. (B) The induction of STPKs in the GoF strains as determined by RNA-seq was within the range observed in historical data (41-45) under physiologic stress conditions for most STPKs. PknK and PknL showed higher induction, most STPKs showed lower induction, suggesting largely physiologic levels of STPK in the GoF strains. (C) STPK abundance is not correlated with the number of DEGs in the GoF strains.

**Figure S6:**
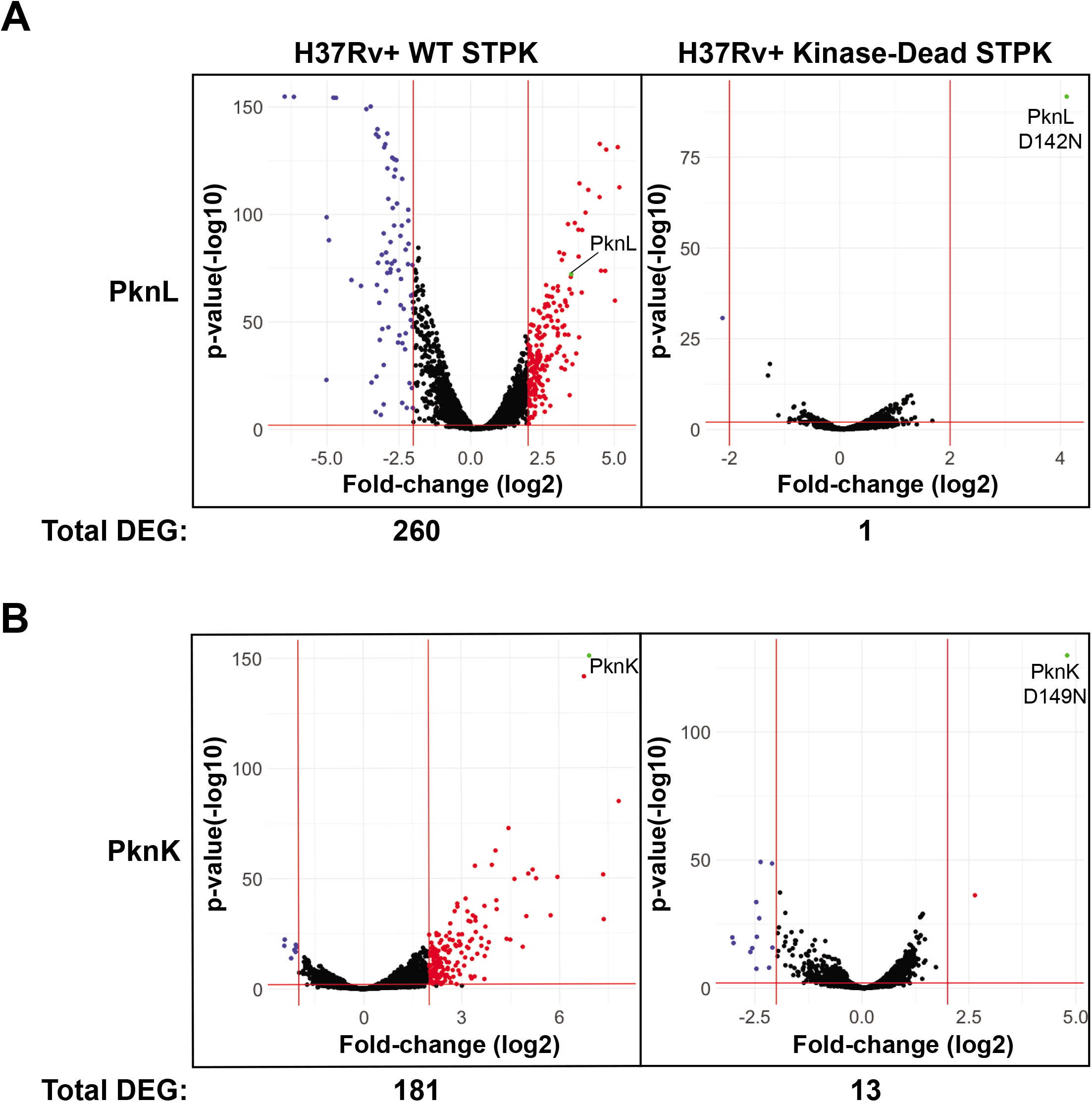
Transcriptional effects of the PknL and PknK kinase dead mutants. (A) The PknL kinase dead mutant reduces the DEGs compared to WT GoF from 260 to 1. PknL WT and kinase dead transcripts are highlighted in green, showing induction in both cases. (B) The PknK kinase dead mutant reduces DEGs compared to WT GoF from 181 to 13, suggesting a small transcriptional effect of the PknK malT domain in the GoF. PknK WT and kinase dead transcripts are highlighted in green, showing induction in both cases.

**Figure S7:**
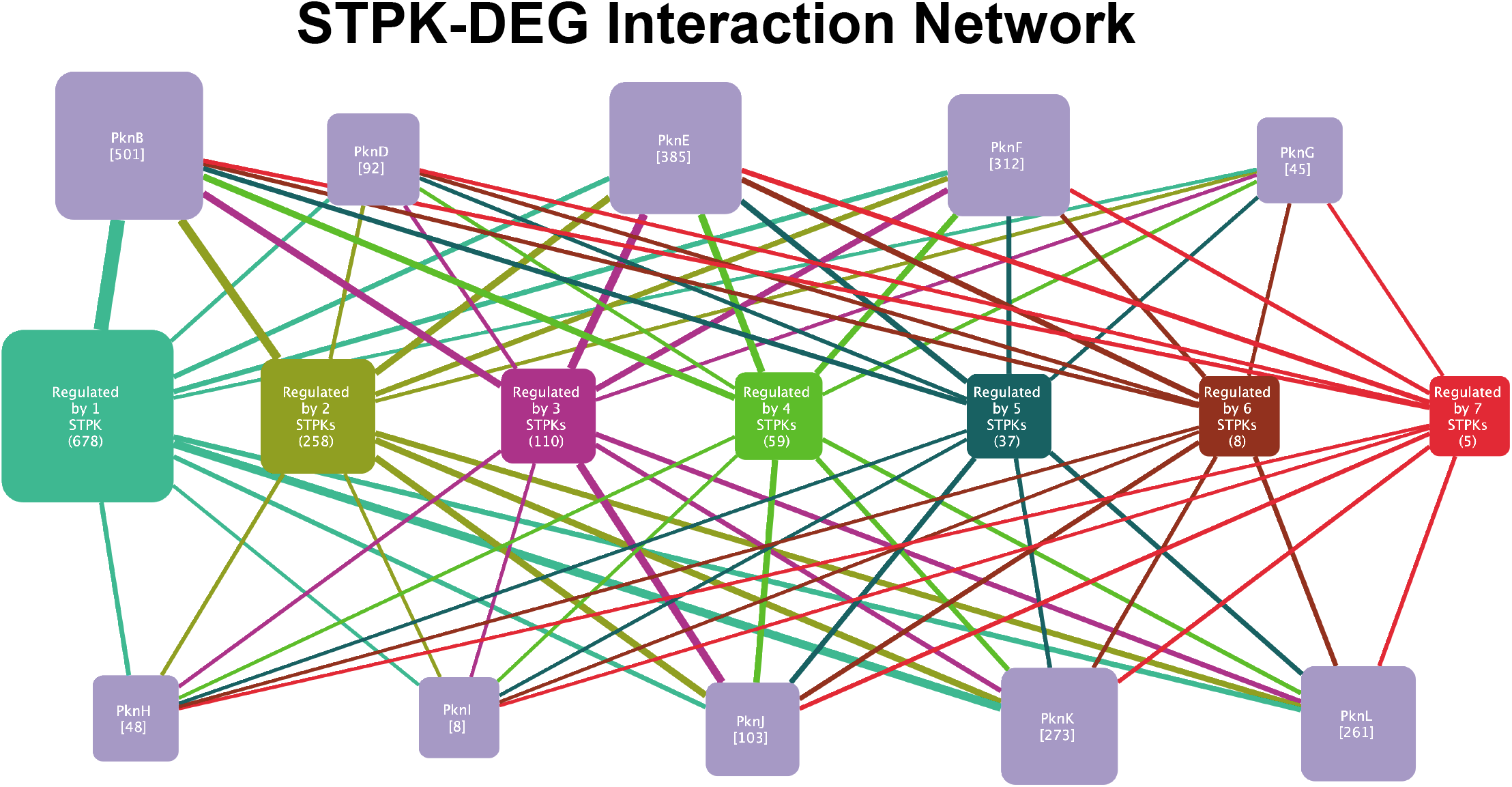
STPK signaling is highly integrated with transcription. STPKs (purple nodes) affect the expression of a large number of genes (number in parentheses). Nodes in the center represent the genes grouped by the number of STPKs they are regulated by. The number in parentheses indicates the number of genes in that group, illustrating the redundancy and/or cooperativity of the network.

**Table S1: Total and unique phosphosites**.

**Table S2: The STPK-related *O*-phosphorylation changes**.

**Table S3: The STPK-related differentially expressed genes**.

## MATERIALS AND METHODS

### *Mtb* STPK mutant strain generation

Loss-of-function (LoF) mutants were constructed by recombineering (22). The recombineering substrate was generated by amplifying 500 bp upstream and downstream of each STPK’s coding sequence by PCR and cloning them 5’ and 3’ of a hygromycin-resistance cassette, respectively. A linear recombineering substrate was amplified by PCR, purified, and transformed into *Mtb* strain H37Rv carrying pNIT-ETc (46). Successful integration was confirmed using DNA sequencing. Gain-of-function (GoF) mutants were constructed using the episomal pDTCF plasmid encoding a tetracycline-inducible promoter, C-terminal FLAG tag, and a hygromycin-selectable marker. STPKs were amplified from *Mtb* strain H37Rv genomic DNA, cloned into the pDTCF plasmid 5’ of the FLAG tag using Gibson assembly cloning, and confirmed with sequencing. Plasmids were transformed into *Mtb* and expression was confirmed by anti-FLAG Western blot. The *pknB* CRISPRi-knockdown strain was constructed using a pJR965 plasmid encoding a tetracycline-inducible dCas9, a kanamycin-selectable marker, and a tetracycline-inducible sgRNA targeting the 5’ coding region of *pknB*. STPK LoF and GoF were confirmed by DNA sequencing and selected reaction monitoring and label-free LC-MS/MS.

### *Mtb* culture

All strains were grown in Middlebrook 7H9 with (10%) OADC and (0.2%) glycerol at 37°C in rolling culture. For experiments involving STPK GoF, cultures were grown in 50 μg/ml hygromycin B. For experiments involving STPK CRISPRi knockdown, cultures were grown in 30 μg/ml kanamycin. WT and STPK LoF strains were grown to stationary phase in triplicate. STPK GoF and CRISPRi knockdown strains were grown in triplicate to OD_600_ 0.4-0.6, induced with 100 ng/ml anhydrotetracycline, and grown to stationary phase.

### Lysate production

*Mtb* cultures were pelleted at 4,000 x g for 5 min at 4°C, washed with phosphate buffered saline, and resuspended in lysis buffer (8 M Urea; 50 mM Tris pH 8.0; 75 mM NaCl; cOmplete protease inhibitor cocktail (Sigma 5892791001); 10 mM sodium fluoride (Sigma 919), 1% phosphatase inhibitor cocktail 2 (Sigma, P 5726), 1% phosphatase inhibitor cocktail 3 (Sigma, P 0044)). Lysates were generated by bead-beating three times at 6 m/s for 30 sec with cooling on ice between steps. Samples were centrifuged at 12,000 x g for 1 min, and the supernatant sterilized by filtration. Protein concentrations were determined by Bradford assay and the lysates stored at −80°C until further processing.

### Proteomic and phosphoproteomic sample processing

1 mg of protein from each sample in 1 ml lysis buffer was incubated for 15 min at 4°C. Protein concentrations were determined by BCA assay (ThermoFisher Scientific). Proteins were reduced with 5 mM dithiothreitol for 1 h at 37 °C, and subsequently alkylated with 10 mM iodoacetamide for 45 min at 25°C in the dark. Samples were diluted 4-fold with 50 mM Tris-HCl, pH 8.0 to decrease a final concentration of urea below 2 M for the digesting with Lys-C (Wako) at 1:50 enzyme-to-substrate ratio. After 2 h of Lys-C digestion at 25 °C, sequencing grade modified trypsin (Promega, V5117) at 1:50 enzyme-to-substrate ratio was added to the samples and samples were incubated at 25°C for 14 h. The reaction was stopped by acidifying the samples with 100% formic acid (Sigma-Aldrich, St. Louis, MO) to a final concentration of 1% formic acid. Acidified samples were centrifuged for 15 min at 1,500 g to clear digest from possible precipitation. Tryptic peptides were desalted on 200-mg tC18 SepPak (Waters, WAT054925) SPE and concentrated down using Speed-Vac concentrator. Final peptide concentration was determined via BCA assay. Desalted peptide samples from each sample were labeled with 10-plex tandem mass tags (TMT, ThermoFisher Scientific) with one of the TMT channels (131) occupied with a pooled mixture of peptides from all the samples, which serves as a reference to normalize across different sets of samples. Peptides (400 μg) were dissolved in 80 μl of 50 mM HEPES, pH 8.5 solution, and mixed with 400 μg of TMT reagent that was dissolved freshly in 20 μl of anhydrous acetonitrile. To quench the labeling reaction, 12 μl of 5% hydroxylamine was added to the samples before for 15 min at RT incubation. Peptides labeled by different TMT reagents were then mixed, dried in Speed-Vac concentrator, reconstituted with 3% acetonitrile, 0.1% formic acid and desalted on tC18 SepPak SPE columns. 3.5 mg of 10-plex TMT-labeled sample was separated on a reversed phase Agilent Zorbax 300 Extend-C18 column (250 mm × 4.6 mm column containing 3.5-μm particles) using Agilent 1200 HPLC System. Solvent A was 4.5 mM ammonium formate, pH 10, 2% acetonitrile and solvent B was 4.5 mM ammonium formate, pH 10, 90% acetonitrile. The flow rate was 1 ml/min and the injection volume was 900 μL. The LC gradient started with a linear increase of solvent B to 16% in 6 min, then linearly increased to 40% B in 60 min, 4 min to 44% B, 5 min to 60% B and another 14 of 60% solvent B. A total of 96 fractions were collected into a 96 well plate throughout the LC gradient. These fractions were concatenated into 6 fractions by combining 16 fractions into one for a total of 6 fractions. For proteome analysis, 5% of each concatenated fraction was dried and re-suspended in 2% acetonitrile, 0.1% formic acid to a peptide concentration of 0.1 mg/ml for LC-MS/MS analysis. The rest of the fractions (95%) were subjected to immobilized metal affinity chromatography (IMAC) for phosphopeptide enrichment (47,48).

### Phosphopeptide enrichment

Fe3+-NTA-agarose beads were freshly prepared using the Ni-NTA Superflow agarose beads (QIAGEN, #30410) for phosphopeptide enrichment. For each of the 12 fractions (∼ 200 ug per each fraction) and for each label-free sample (100 ug), peptides were reconstituted to 0.5 μg/μl in IMAC binding/wash buffer (80% acetonitrile, 0.1% trifluoroacetic acid) and incubated with 10 μl of the Fe3+-NTA-agarose beads for 30 min at RT. After incubation, the beads were washed 2 times each with 50 μl of wash buffer and once with 50 μl of 1% formic acid on the stage tip packed with 2 discs of Empore C18 material (Empore Octadecyl C18, 47 mm; CDS Analytical, 98-0604-0217-3). Phosphopeptides were eluted from the beads on C18 using 70 μl of Elution Buffer (500 mM potassium phosphate buffer). 50% acetonitrile, 0.1% formic acid was used for elution of phosphopeptides from the C18 stage tips. Samples were dried using Speed-Vac, and later reconstitute with 12 μl of 3% acetonitrile, 0.1% formic acid containing 0.01% DDM (n-Dodecyl-beta-Maltoside) for LC-MS/MS analysis.

### Global and phosphoproteome analysis

Both TMT and label-free global and phosphopeptide enriched samples were subjected to a custom high mass accuracy LC-MS/MS system as previously described (11), where the LC component consisted of automated reversed-phase columns prepared in-house by slurry packing 3-μm Jupiter C18 (Phenomenex) into 35-cm x 360 μm o.d. x 75 μm i.d fused silica (Polymicro Technologies Inc.). The MS component consisted of a Q Exactive HF Hybrid Quadrupole-Orbitrap mass spectrometer (Thermo Scientific) outfitted with a custom electrospray ionization interface. Electrospray emitters were custom made using 360 μm o.d. × 20 μm i.d. chemically etched fused silica capillary. Analysis of the phosphoproteome samples applied similar conditions as used in the global proteome sample analysis, except that the spray voltage was 2.2 kV. All other instrument conditions were set as previously described (49). Raw spectral data and analysis information is available via the MassIVE database for the public accession #MSV000088254. LC–MS/MS raw data were converted into dta files using Bioworks Cluster 3.2 (Thermo Fisher Scientific, Cambridge, MA, USA). The MSGF+ algorithm was used to search MS/MS spectra against the *Mtb* database (RefSeq H37Rv_uid57777_2014-08-14, 20198 entries). Search parameters included 20 ppm tolerance for precursor ion masses, +2.5 Da and −1.5 Da window on fragment ion mass tolerances, no limit on missed cleavages, partial tryptic search, no exclusion of contaminants, dynamic oxidation of methionine (15.9949 Da), static IAA alkylation on cysteine (57.0215 Da), and static TMT modification of lysine and N-termini (+144.1021 Da) for TMT analyses. The decoy database searching methodology (50,51) was used to control the false discovery rate at the unique peptide level to <0.01% and subsequent protein level to <0.1% (52). Quantification for TMT was based upon initially summing to the peptide (phospho) or protein (global) level the sample specific peptide reporter ion intensities captured for each channel across all 24 or 12 analytical fractions. Final data for statistical analysis was the scaling and central tendency normalization of each peptide or protein summed value across all observations within each TMT10 experiment to adjust for experiment specific variability. Label-free quantification used MaxQuant (53) generated LFQ values at the peptide (phospho) and protein (global) levels as previously described (54). All collected protein and peptide abundance values were subjected to statistical comparison using WT as the basis for comparison. Primary comparisons were based upon ANOVA analysis at a <0.005 p-value threshold coupled with a fold change criteria minimum of +/-2 between abundances.

### Selected reaction monitoring

SRM was performed on the panel of WT, GoF, and LoF mutants as described above. Crude heavy peptides labeled with ^13^C/^15^N on C-terminal lysine and arginine were purchased from vivitide (Gardner, MA). Trypsin digested samples that had been stored at −80°C until use were processed as previously described (55). For each sample the digested peptides were diluted to 0.25 μg/μl containing standards at a final concentration of 20 fmol/μl. All the samples were analyzed with a nanoACQUITY UPLC system (Waters Cooperation, Milford, MA) coupled online to a TSQ Altis triple quadrupole mass spectrometer (Thermo Fisher Scientific, San Jose, CA). The LC-SRM platform was configured and utilized as previously described (56). The abundance of PknE and PknI in wild-type, PknE LOF mutant, and PknI LOF mutant were too low to be detectable through the standard LC-SRM analysis. An extra PRISM procedure was applied to these low abundance samples to enrich the targeted peptides for PknE and PknI before LC-SRM analysis (57). SRM data acquired on the TSQ Altis were analyzed using Skyline-daily software (version 21.0.9.139) (58). Peak detection and integration were determined based on retention time and the relative SRM peak intensity ratios across multiple transitions between light peptide and heavy peptide standards (59). All the data were manually inspected to ensure correct peak assignment and peak boundaries. The peak area ratios of endogenous light peptides and their heavy isotope-labeled internal standards (i.e., L/H peak area ratios) were then automatically calculated by Skyline-daily, and the average peak area ratios from all the transitions were used for quantitative analysis of the samples. For targets that had more than one surrogate peptide, correlation graphs were plotted to verify a strong correlation and ultimately the peptide that had the most sensitive response was selected for obtaining quantitative values.

### RNA Isolation and RNA sequencing

*Mtb* cultures were pelleted at 4000 × g for 5 min at 4°C, resuspended in Trizol, transferred to a tube containing Lysing Matrix B, and lysed via bead beating using 30 sec shaking at 6 m/s in a homogenizer three times, with cooling on ice. The samples were centrifuged at 21,000 x g for 1 minute, and the supernatant was transferred to a tube containing 300 μl chloroform and Heavy Phase Lock Gel. The tubes were inverted for 2 minutes and centrifuged at 21,000 x g for 5 minutes. RNA in the aqueous phase was precipitated using 300 μl isopropanol and 300 μl high-salt solution (0.8 M Na citrate, 1.2 M NaCl). RNA was purified using a RNeasy kit with one on-column DNase treatment. Total RNA was quantified using a Nanodrop. Messenger RNA was enriched by first depleting ribosomal RNA from samples using the Ribo-Zero rRNA removal (bacteria) magnetic kit (Illumina). The purified mRNA was prepared for Illumina sequencing using the NEBNext Ultra RNA Library Prep Kit for Illumina according to the manufacturer’s instructions in combination with AMPure XP reagent for clean-up of adaptor-ligated DNA. Each replicate was barcoded in the DNA library using the NEBNext Multiplex Oligos for Illumina (Dual Index Primers Set 1). 30-40 libraries were multiplexed per sequencing run, and libraries were quantified using the KAPA qPCR quantification kit. The libraries were sequenced at the University of Washington Northwest Genomics Center with the Illumina NextSeq 500 High Output v2 Kit. Read alignment was performed using the custom processing pipeline that uses Bowtie 2 utilities, available at https://github.com/robertdouglasmorrison/DuffyNGS and https://github.com/robertdouglasmorrison/DuffyTools.

### Phosphorylation motif analysis

The custom background peptideome was generated by using the *Mtb* H37Rv proteome from UniProt.org (Proteome ID: UP000001584, modification date 12/1/2019). A custom R script was used to partition the proteome into unique, 11 amino acid long peptides. Sequences were further subdivided into lists whose central residue was either serine, threonine, or tyrosine. The foreground peptidome was generated by compiling the differentially modulated phosphosites for each kinase mutant. An R script was used to create a 11 amino acid sequence centered on the modulated phosphosite. Additionally, the same script was used to generate a list of all phosphosites identified in the WT phosphoproteome (basal phosphoproteome). For the motif analysis, four separate analyses were performed for each of the kinase mutants and the basal phosphoproteome: a site-centered or a Ser/Thr/Tyr-centered motif search. Foreground and respective background peptidomes were uploaded to the pLogo web application for analysis using default parameters.

### *In vitro* phosphorylation assay

The kinase domains of PknB-PknL were cloned from *Mtb* genomic DNA into inducible pET28b protein expression plasmid using Gibson assembly cloning and confirmed by DNA sequencing. pET28b plasmids were transformed into *E. coli* BL21 for protein expression. Cultures were grown to 0.4-0.6 OD, induced with 250 μM IPTG, and grown at 20°C for 18-24 hours. Cultures were pelleted at 4000 x g for 20 minutes, resuspended with lysis buffer (20 mM Tris-HCl pH 7.5, 150 mM NaCl, 20 mM imidazole), and lysed by sonication. Lysates were centrifuged to remove the insoluble fraction, and purified using affinity chromatography with Ni-NTA. Proteins were further purified using size exclusion chromatography and stored in size exclusion buffer (20 mM Tris-HCl, 150 mM NaCl, 5% glycerol, pH 7.4). For the auto-phosphorylation assay, 1 μM STPK was combined with reaction buffer (50 μM MnCl_2_, 50 μM MgCl_2_, 1 mM DTT, 50 mM Tris pH 8.0, 75 mM NaCl) and initiated with 50 μM ATP and 15 μCi ATP [γ-32P]. The reaction was incubated at 37°C for 30 min. To terminate the reaction, 4x loading buffer with β-mercaptoethanol was added and incubated at 95°C for 5 minutes. Proteins were separated by SDS-PAGE in MES buffer and 4-15% Bis-Tris Plus gels. Gels were dried using a gel dryer at 80°C for 3 hours. Phosphor screens were exposed to the dried gels overnight and then imaged using a Sapphire biomolecular imager. For Zur phosphorylation assay, 10 μM Rv2359 was combined with 1 μM STPK in reaction buffer (50 μM MnCl_2_, 50 μM MgCl_2_, 1 mM DTT, 50 mM Tris pH 8.0, 75 mM NaCl), with 50 μM ATP and 15 μCi ATP [γ −32P], and then incubated at 37°C for 30 min. Proteins were separated by SDS-PAGE and visualized as above.

### Zur mutagenesis

Zur phosphomimetic (Thr67Asp) and phosphoablative (Thr67Ala) mutants were generated by QuikChange site-directed mutagenesis. In summary, overlapping primers containing the desired mutation were used to linearize and introduce the mutation into the pET28b plasmid containing Zur. PCR products were gel extracted and DpnI-digested, followed by transformation into competent *E. coli*. Mutagenesis was confirmed by DNA sequencing.

### Electrophoretic Mobility Shift Assay (EMSA)

Cy5-labeled DNA probe was made by resuspending three oligos (Integrated DNA Technologies) to 100 μM in dsDNA annealing buffer (10 mM Tris-HCl-pH 7.5, 100 mM NaCl, 1 mM EDTA). Oligo 1 corresponded to 100-200 bp binding region of Rv2359. Oligo 2 corresponded to the reverse complement of Oligo 1, as well as a 12-nucleotide sequence at the 3’ end to which Oligo 3 was the reverse complement. Oligo 3 was a 12-mer with Cy5 covalently coupled to the 5’ end. The three oligos were combined to a final concentration of 100 μM, vortexed, and heated to 95°C for 10 min. The reaction was cooled to room temperature over 3-4 hours protected from light and used for EMSA. For the EMSA, various concentrations of purified recombinant Rv2359 WT or mutant protein were combined with 2 μM final DNA probe in reaction buffer (20 mM Tris-HCl pH 8.0, 75 mM NaCl, 1 mM DTT, 50 μM MgCl_2_, 50 μM MnCl_2_, 5% glycerol, 1% zinc sulfate, 50 μg/mL bovine serum albumin, 50 μg/mL salmon sperm DNA) and incubated for 30 min at room temperature, protected from light. After incubation, 4 μl native sample loading buffer was added to each sample. The whole reaction was loaded on a native 10% Tris-Glycine gel, and gel was maintained on ice and ran at a constant 100 volts for 2 hours. The gel was rinsed with water and visualized on a Li-cor Odyssey scanner. Bands were quantified using the Odyssey software.

## ACKNOWLEDGEMENTS

This work was supported by NIH grants R01AI117023, R21AI137571, and R03AI131223 and by a grant by the American Lung Association to C.G. A.F. was supported by the Interdisciplinary Program in Bacterial Pathogenesis 5T32AI053396. Portions of this research were supported by NIH NIGSM GM103493. Some work was performed in the Environmental Molecular Sciences Laboratory, a U.S. Department of Energy Office of Biological and Environmental Research national scientific user facility located at Pacific Northwest National Laboratory in Richland, Washington. Pacific Northwest National Laboratory is operated by Battelle for the U.S. Department of Energy under Contract No. DE-AC05-76RLO 1830.

